# Single-cell transcriptomics reveals that glial cells integrate homeostatic and circadian processes to drive sleep-wake cycles

**DOI:** 10.1101/2023.03.22.533150

**Authors:** Joana Dopp, Antonio Ortega, Kristofer Davie, Suresh Poovathingal, El-Sayed Baz, Sha Liu

## Abstract

The sleep-wake cycle is determined by circadian and sleep homeostatic processes. However, the molecular impact of these processes and their interaction in different brain cell populations remain unknown. To fill this gap, we profiled the single-cell transcriptome of adult *Drosophila* brains across the sleep-wake cycle and four circadian times. We show cell type-specific transcriptomic changes with glia displaying the largest variation. Glia are also among the few cell types whose gene expression correlates with both sleep homeostat and circadian clock. The sleep-wake cycle and sleep drive level affect expression of clock gene regulators in glia, while diminishing the circadian clock specifically in glia impairs homeostatic sleep rebound after sleep deprivation. These findings reveal a comprehensive view of the effects of sleep homeostatic and circadian processes on distinct cell types in an entire animal brain and reveal glia as an interaction site of these two processes to determine sleep-wake dynamics.

## Main

Sleep is regulated by two independent intrinsic processes: the circadian and sleep homeostatic system^1^. The circadian rhythm primarily regulates the timing of sleep, known as Process C. It is generated by the core molecular oscillator, which comprises a transcriptional-translational feedback loop consisting of core clock genes^2^. At the cellular level, our current understanding of the circadian timing system focuses on the circadian pacemaker regions/neurons, such as the suprachiasmatic nuclei (SCN) in mammals^3^ and 150 clock neurons in *Drosophila*^4^. However, it remains unclear whether and how Process C affects the transcriptomes of any given brain cell population apart from pacemaker regions/neurons.

The sleep homeostat monitors the sleep need that accumulates with the amount of time that an animal has been awake to determine the sleep drive, known as Process S. In contrast to the well-defined molecular and cellular mechanisms of the circadian clock, our understanding of the nature of the sleep homeostat remains limited. Previously, the effects of the sleep-wake cycle and sleep homeostasis on the transcriptome were studied in bulk samples, either from brain tissue or bulk synaptosomes^5–10^ in mammals, or whole brain tissue in flies^7, 8^. However, these studies are inconsistent, possibly due to averaging transcriptomic changes of heterogeneous cell populations (**Extended Data Fig. 1**). Recently, single-cell RNAseq has been applied to study the transcriptional changes during sleep homeostasis in mice^11, 12^.

However, these studies only focused on certain regions of the brain. Therefore, similar to circadian cycling transcripts, we are missing a comprehensive and unbiased understanding of sleep/wakefulness and sleep homeostasis-associated transcriptomic changes across cell populations of an entire brain.

Here, we sampled adult *Drosophila* brains at different sleep, wakefulness, and sleep pressure states at different circadian times, and performed single-cell RNA sequencing, thereby creating a comprehensive transcriptional atlas of the sleeping animal brain (see https://www.flysleeplab.com/scsleepbrain for the single-cell gene expression atlas and analyses links). We find that sleep/wakefulness states, sleep homeostasis, and circadian rhythm have different transcriptional correlates depending on the cell identity. Interestingly, our data also suggest that gene expression in most cell populations correlates either with Process C or S, with the exception of glial cells. Glia instead are affected by both processes simultaneously, leading us to suggest a model whereby homeostatic and circadian processes directly interact at glial cells to regulate sleep.

## Results

**Single-cell transcriptomes of fly brain cells at different sleep/wakefulness states and circadian times.**

We performed single-cell RNA sequencing with 10x droplet microfluidics on adult *Drosophila* central brains (without optic lobes), sampled at distinct points of the sleep-wake cycle across different circadian times (**Fig. 1a**). These sample points can be 1) grouped by four Zeitgeber (ZT) times to analyse the transcriptional correlates of circadian rhythms; 2) grouped by “sleep” and “wakefulness” states according to the animal’s vigilance status at the time of sample collection to examine “sleep/wakefulness” correlates; and 3) selected and ordered by the fly’s level of sleep drive to determine molecular changes associated with the sleep homeostat. To minimize technical batch effects that may mask true biological responses, we applied demultiplexing based on natural variation between wild-type genotypes^13, 14^. Specifically, instead of associating each batch to a different sleep or wakefulness state, we associated a different *Drosophila* Genetic Reference Panel (DGRP)^15^ line to each behavioural condition (**Fig. 1b**, **Supplementary Table 1**). The full genome of all DGRP lines has been sequenced^16^, thus they can be distinguished from one another by their unique single nucleotide polymorphisms^15^. Thanks to this natural genetic variation, the RNA sequencing data discloses the associated sleep or wakefulness states, which allows the pooling of brains from different behavioural conditions in a single batch (**Fig. 1b**) (see also Methods). In addition to minimizing technical batch effects, we also counteracted potential genotype-specific effects by repeating the same conditions with different DGRP lines with similar sleep profiles in subsequent batches (**Supplementary Table 1**). To find the DGRP lines with the most similar sleep behaviour, we carefully chose them by 1) pre-selecting thirty-six DGRP lines based on sleep architecture metrics previously reported in a large collection of DGRP lines^17^ and 2) thoroughly screening those thirty-six lines across various sleep parameters. The criteria for selection were (1) a robust amount of consolidated night-time sleep and a low amount of day-time sleep, and (2) low between and within genotype variability (**Extended Data Fig. 2a**). Principal component (PC) analysis of these sleep parameters showed that the ten selected lines group together closer than the discarded lines (**Extended Data Fig. 2b**). Similar clustering of selected vs. discarded lines is also visible when plotting the probabilities to transition from a wake to a sleep state (pDoze) or vice versa (pWake), which indirectly assess sleep drive and sleep depth in *Drosophila*^18^ (**Extended Data Fig. 2c-d**). The low variability between the sleep phenotype of the selected ten DGRP lines allowed us to repeat each condition with several lines in the same and in different batches (**Supplementary Table 1**). The success of this strategy is attested by the homogenous distribution of cells from different genotypes (**Extended Data Fig. 3a**), that blend within cell subtypes even without applying batch integration algorithms (**Extended Data Fig. 3b**).

**Figure 1.**
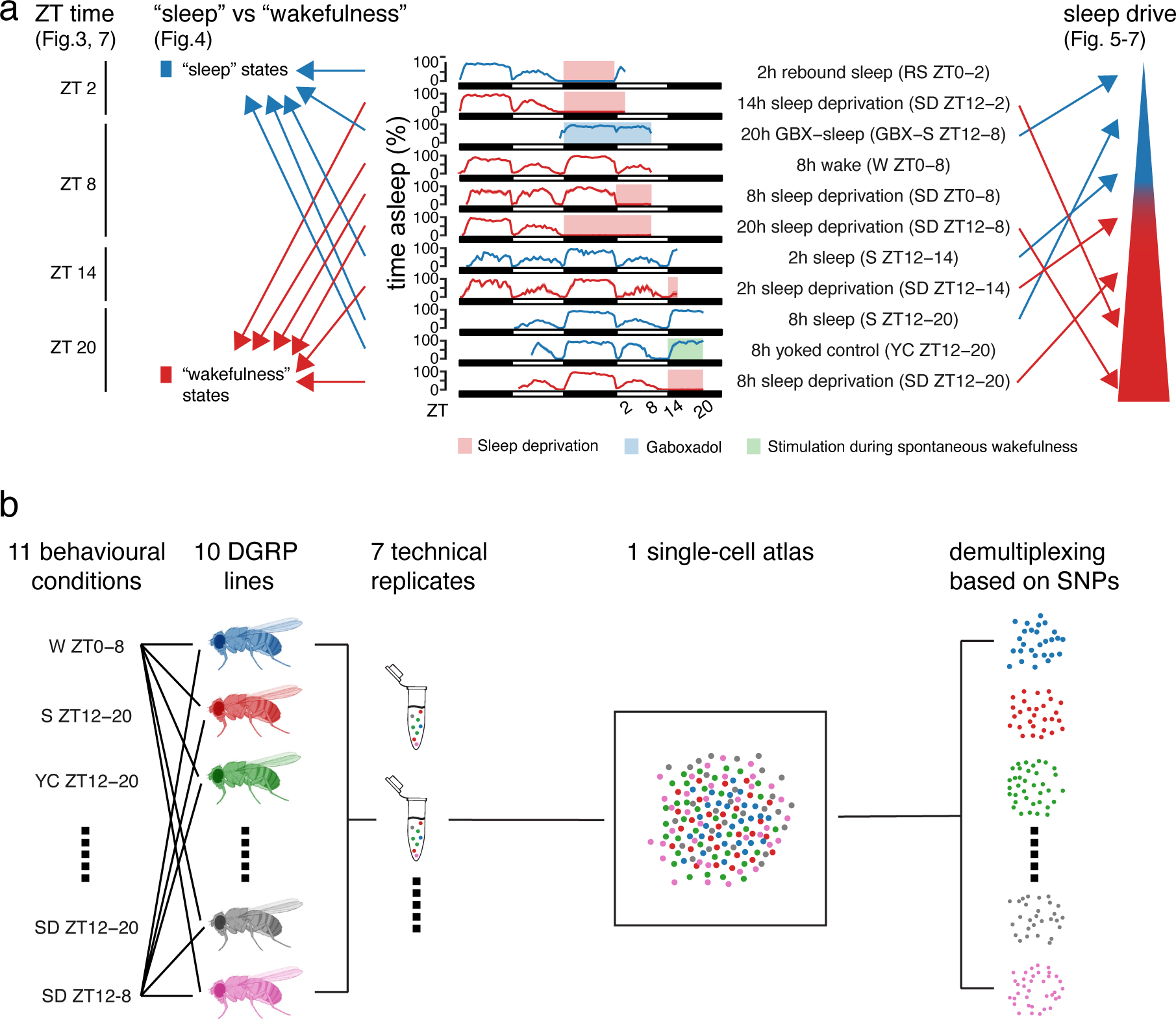
Sampling flies at sleep and wakefulness states for subsequent transcriptional profiling. **a.** Flies are sampled at four different Zeitgeber (ZT) times and eleven different sleep or wakefulness states. Seven of the conditions are ordered by accumulated sleep pressure. The corresponding downstream analyses of each of the three correlates are described in the corresponding fgure displayed in parentheses. **b.** In each of seven technical replicates (runs), each condition was linked to two or three DGRP lines. The link between condition and DGRP line was changed in every run. The lies’ central brains were dissected and the tissue dissociated in a single tube minimizing batch effects. Conditions were separated by demultiplexing the sequenced reads based on unique SNPs of DGRP lines.

To obtain an overview of the captured cell types, we first performed dimensionality reduction on all 106,762 cells combined, and identified 214 clusters of cells (**Fig. 2a**). We annotated cells based on previously used marker genes in the fly brain cell atlas^19, 20^, allowing us to successfully assign 22,988 cells to one of 25 known cell types (21.5%) (**Fig. 2b-c, Supplementary Table 2**), including five glial subtypes, Kenyon Cells (KCs), clock neurons, and cell types containing known sleep/wakefulness regulating circuits such as, non-PAM dopaminergic neurons (DANs)^21–23^, Tyraminergic/Octopaminergic neurons^24, 25^, and ellipsoid body (EB) ring neurons^26–28^. Another major sleep-regulating cell type, the dorsal fan-shaped body (dFB) neurons was annotated by correlating the previously published transcriptome of FAC-sorted dFB neurons with our gene expression data (**Extended Data Fig. 4**).

**Figure 2.**
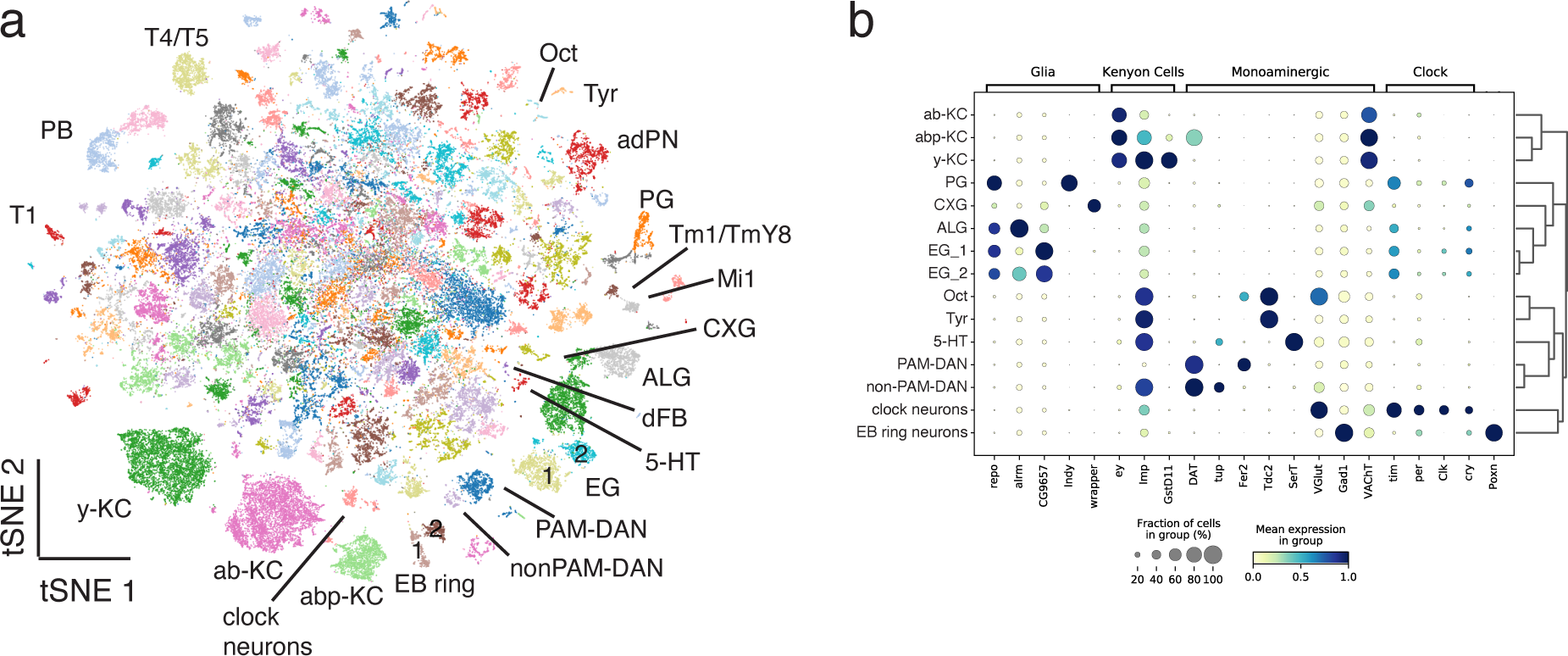
Cell type annotations in a single-cell atlas of the sleeping fruit fly. **a.** tSNE plot of entire dataset of 106.762 single cells with annotated clusters and expression of key marker genes of glial and neuronal cell types used to annotate clusters. KC = Kenyon Cells, EG = ensheathing glia, ALG = astrocyte-like glia, PG = perineurial glia, CXG = cortex glia, Oct = Octopaminergic, Tyr = Tyraminergic, 5-HT = Serotonergic, PAM-DAN = protocerebral anterior medial part of dopaminergic neurons, EB = ellipsoid body, adPN = andero-dorsal projection neurons, PB = protocerebral bridge neurons, dFB = dorsal fan-shaped body. **b.** Marker genes expression across most clusters annotated in **a.**

### Cycling of core circadian genes in clock neurons and glia

The expression levels of circadian clock gene transcripts cycle in a daily manner in clock neurons^2^. To validate our single-cell transcriptomic dataset, we asked whether the data accurately capture cycling expression of core clock genes between the four sampled ZT time points. Indeed, as previously reported in clock neurons^29, 30^, *period* (*per*) and *timeless* (*tim*) transcripts are expressed at higher levels in the early night compared to the early day, while the opposite applies to *cryptochrome* (*cry*) and *Clock* (*Clk*) mRNA (**Fig. 3a**). Beyond validating our dataset, we examined whether and how the clock genes are cycling in all remaining cell populations. In mammals, cycling of clock gene expression in cell populations other than the SCN has been reported, for example in cortical regions^31^. Interestingly, we report that the expression and cycling of these genes is restricted specifically to clock neurons and glial cells (**Fig. 3a-b, Extended Data Fig. 5a-e**). To confirm this, we tested the activity of the *Clk* regulatory network (regulon) across all cells, by applying SCENIC^32^. This regulon is defined by the previously identified *Clk* binding element E-box sequence and its target genes, including *tim*, *cry* and *vrille* (*vri*)^33^. Our data shows that the *Clk* regulon is only active in clock neurons and most glial subtypes, not in other neurons (**Fig. 3c**), further indicating that core clock genes are expressed and cycle specifically in *Drosophila* clock neurons and glia.

**Figure 3.**
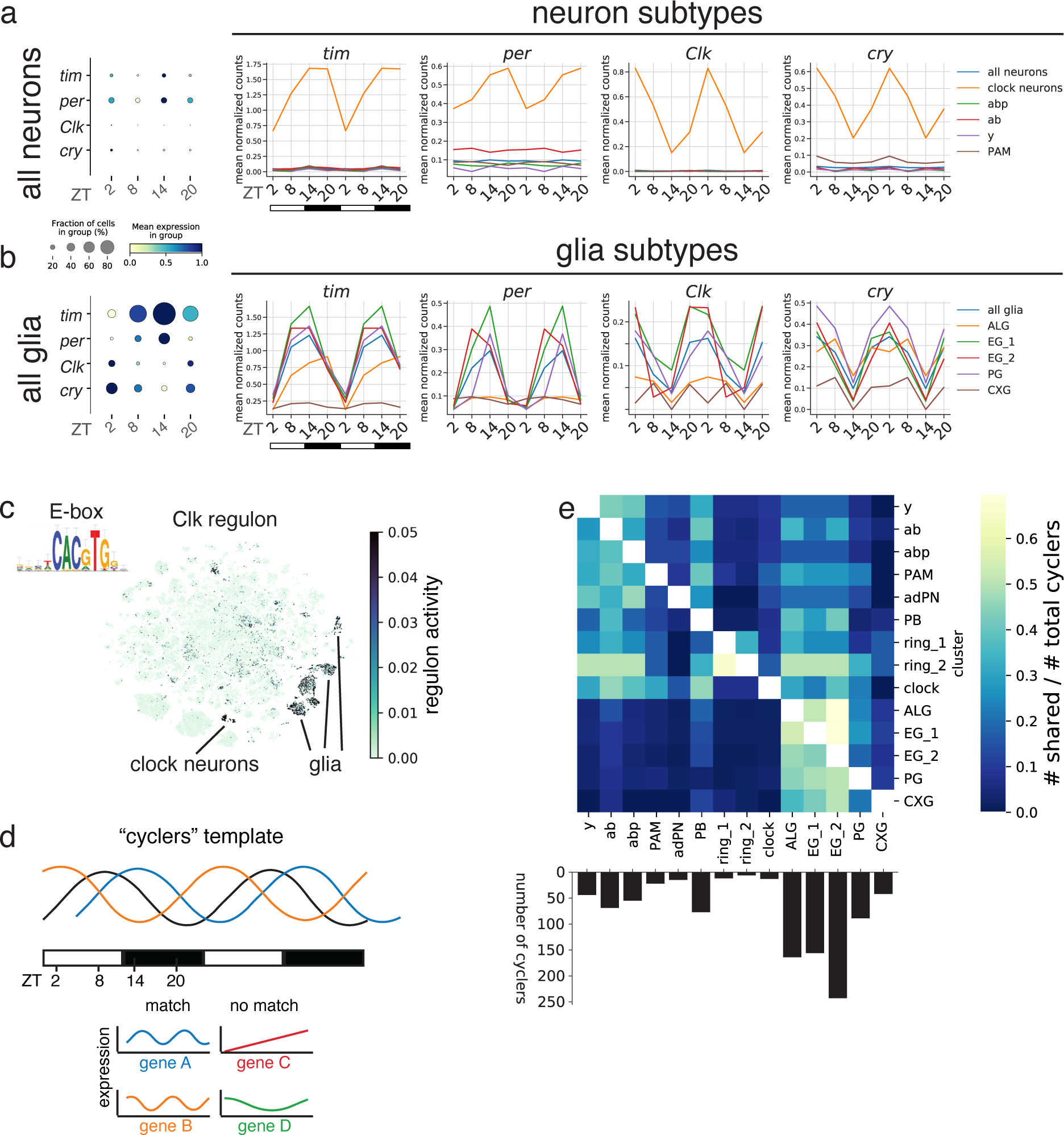
Oscillating transcripts in neurons and glia. **a-b.** Circadian expression levels of core clock genes per, tim, cry and Clk averaged across all neurons, some neuronal subtypes (a) and all glia and glial subtypes (b). Size of dot indicates fraction of cells in group. Mean expression in group is normalized to expression of gene across the four ZT time points. Data in line plot is plotted twice to better visualize cycling patterns. **c.** Clk regulon and its activity across all cell types. **d.** Schematic of template to detect cycling transcripts. **e.** Heatmap of intersecting cycling genes between all annotated cell types with at least one shared gene relative to the total number of cycling genes (at least five genes, bottom bar plot) by cell type.

Fly glia have previously been associated with expression of clock genes, especially astrocytes (ALG)^34^ and perineurial glia (PG)^35^. Interestingly, our data show that the molecular clock runs with different phases depending on the cell type. Specifically, the expression of *tim* and *per* mRNA is high throughout the night in clock neurons, while in PG and ensheathing glia (EG), its expression decreased already at ZT 20 (**Fig. 3b, Extended Data Fig. 5a-e**). In contrast, *tim* expression in astrocytes only peaks at ZT 20 in astrocytes. The delayed clock in astrocytes compared to neurons is also observed in the mouse suprachiasmatic nucleus (SCN)^36^. Taken together, the expression of key clock genes in glia in addition to clock neurons, suggests that these cells are directly involved in circadian regulation of rhythmic behaviours, including sleep.

### Circadian “cyclers” are more enriched in glial than neuronal cell types

Next, we comprehensively identified all circadian oscillating transcripts (cyclers) in all cell types by applying the JTK cycle algorithm (**Fig. 3d**, see also Methods)^38^. While the expression of the molecular clock is restricted to clock neurons and glial cells, the majority of cell clusters (82.5%) have at least one cycler (**Extended Data Fig. 5f**) and 14 of the 19 annotated clusters have at least five cyclers (**Fig. 3e**). Similar to the analysis of core clock genes, glial cells stand out, this time by showing the highest number of cyclers, especially considering they typically express less genes than neurons (**Extended Data Fig. 5g**). When comparing the overlap of cyclers between all cell types, we find that only between 20% and a maximum of 50% of cyclers are shared. We identify a higher number of shared cyclers between closely related cell types, such as the three KC subtypes and particularly between neuropil-associated glial subtypes (ALG and EG) (**Fig. 3e**). These data suggest different cell populations respond differently to Process C, with glial cells being most affected.

### Sleep/wakefulness correlates differ between cell populations

Next, we asked whether we can detect transcript level changes across sleep/wakefulness states (referred to as sleep/wakefulness correlates), similarly to circadian times. We assigned each condition to a “sleep” or “wakefulness” group based on whether the animal had consolidated sleep or wake prior to sampling (**Fig. 1a, and 4a,** also see Methods). The “sleep” group includes spontaneous sleep, recovery sleep following sleep deprivation, yoked control, and Gaboxadol (GBX) induced sleep^39^. We grouped spontaneous wake and forced wake (sleep deprivation) in order to regress out unwanted effects of circadian time and potential stress induced by mechanical sleep deprivation. Then, we performed differential expression analysis between the two groups for each cell population separately and asked whether transcriptomic changes between sleep and wakefulness states differ between distinct cell populations. We found that 46.7% of all clusters and 57.9% of annotated clusters have sleep/wakefulness correlates (**Fig. 4b, Extended Data Fig. 6a**). KCs and glia have the highest number of differentially expressed genes (DEGs) among the annotated cell types.

**Figure 4.**
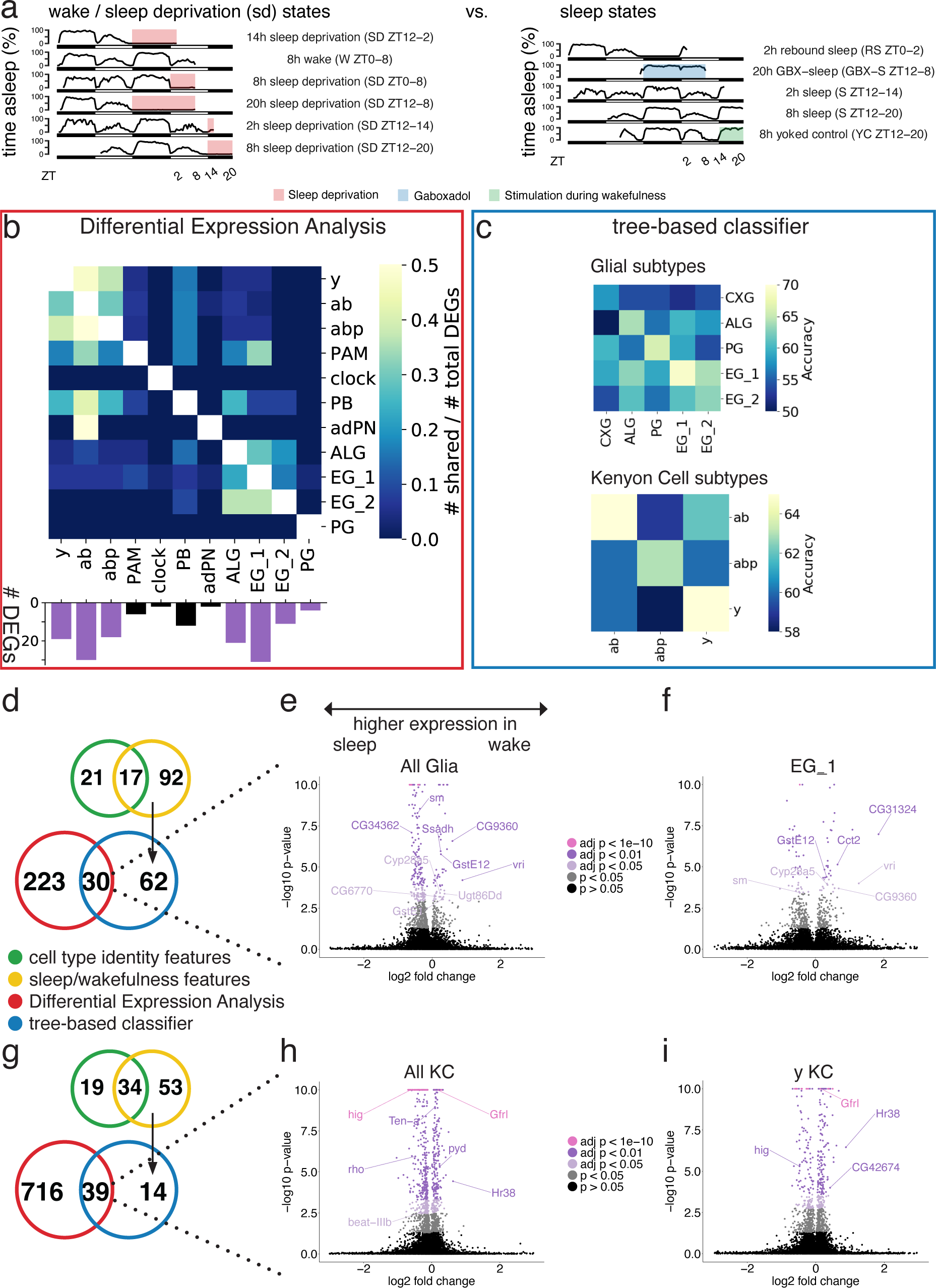
Transcriptomes of glia and KCs change differently between sleep and wake. **a.** Grouping of wake / sleep deprivation states and sleep states to compare them in differential expression analysis and a tree-based classifier. **b.** Heatmap of intersecting differentially expressed genes (DEGs) between all annotated cell types with at least one shared gene relative to the total number of DEGs by cluster. The bottom bar plot shows the amount of total DEGs with adj. p-value < 0.05 and log foldchange <-0.5 or >0.5. **c.** Classifying glia (top) and KC (bottom) subtypes into sleep or wake label reveals that performance is highest for the same subtype. **d-i.** For glia (d) and KC (g), candidate genes were filtered by (1) removing cell type features from sleep/wake features identified by separately trained EBMs and (2) overlapping significant DEA and EBM results. Volcano plots highlighting a selection of the common genes between the two methods for all glia (e), EG_1 (f), all KC (h) and y-KC (i).

Considering that glia express the lowest number of genes among all cell types (**Extended Data Fig. 6b**), these cells may be even more affected by sleep/wakefulness relative to their total expressed genes compared to neurons. Interestingly, the majority of identified DEGs in each of the annotated cell types is unique to that cell type (**Fig. 4b**). On average only 12.8% or 13.5% of DEGs are shared within neuronal or glial subtypes, respectively. Similar to the cyclers, the overlap of sleep/wakefulness correlating transcripts is even smaller between neurons and glia, averaging between 3-4% only, with the exception of PAM dopaminergic neurons and protocerebral bridge neurons that share between 17-33% of their sleep/wakefulness correlating genes with ALG and EG_1. This low overlap between closely related cell subtypes suggests that sleep-wake cycles affect their transcriptome in unique ways.

To further probe the differences between the closely related KC and glia subtypes, we applied a complementary, independent approach to differential expression analysis. We asked whether a tree-based classifier can learn the transcriptome make-up of one cell subtype during sleep and wakefulness states and subsequently, how accurately this model performs in classifying cells of another closely related subtype into either of these states (**Fig. 4c**). This approach was applied to KC and glial cell subtypes separately. To ensure that the classifier does discriminate between sleep and wakefulness states rather than cell identity, we excluded marker genes between either KC or glial subtypes up to the point that the subtypes merged into one another in the two-dimensional UMAP space (**Extended Data Fig. 7a, b**). The accuracy of the classifier is illustrated in a confusion matrix, where the colour corresponds to the probability to accurately assign a “sleep” or “wakefulness” label per cell type. We found that the classifier performs better for the cell subtype it was trained on, than on other related ones (**Fig. 4c**). To ensure that the classifier’s improved performance is not driven by overfitting to the cell type (i.e., learning cell identity features), we randomly shuffled the sleep and wake label of cells within their subtype identity. Then, the classifier cannot distinguish between the sleep and wake states even in the same subtype, suggesting that the classifier truly learns sleep/wakefulness, rather than cell identity features (**Extended Data Fig. 7c, d**). Taken together, both the differential expression analysis and classification approach suggest that depending on their cell identity, different cell populations have different sleep/wakefulness correlates.

Next, we asked what kind of genes are altered between sleep and wakefulness in glia and KCs. To gain confidence in the candidate gene list, we first trained another classifier to detect cell identity features between glial and KC subtypes. Then, we subtracted those cell identity features from those used by the sleep/wakefulness classifier (**Fig. 4d, g**). To reduce the false positive rate further, we considered only genes as significantly deregulated if they remained after the filtering step above and if they were identified as significant in the differential expression analysis, leaving 30 and 39 sleep/wakefulness correlating transcripts in glia and KCs, respectively (**Fig. 4d, g**). Among the filtered 39 sleep/wakefulness correlates of KCs we find Hr38, an activity-dependent gene in insects^40, 41^. To validate its increased expression during sleep deprivation and wakefulness compared to sleep in KCs specifically, we performed fluorescent *in situ* hybridization during sleep and sleep deprivation. In accordance with our single cell data, Hr38 mRNA in KCs increases expression after sleep deprivation (**Extended Data Fig. 8**). The sleep/wakefulness correlates in glia include metabolism related genes (*CG9360*, *Cyp28a5, GstE12, GstE1, mdh1, Ssadh, and Ugt86Dd*), genes involved in protein synthesis and homeostasis (*Rpl41, Cct2, CG34362, CG6770, sm*), and genes regulating core circadian clock (*CG31324* and *vri*). In contrast, sleep/wakefulness correlates in KCs include many genes involved in axon and synapse development and function: *Ten-a, rho, pyd, hig, Gfrl, CG42674, Hasp, beat-IIIb*.

### Template matching successfully captures known cell types involved in Process S

While the sleep/wakefulness correlates detect how the transcriptome changes in a binary manner, we next assessed whether it also changes gradually with the level of sleep pressure (sleep drive correlates). We have sampled flies at multiple sleep and sleep deprivation states that can be ordered by their level of sleep drive, from 20 h of GBX-induced sleep to 20 h of sleep deprivation (**Fig. 5a**). To detect transcripts whose expression correlates with the gradual increase of sleep drive, we adopted a feature selection method^42^. Similar to the JTK algorithm for identification of cyclers which calculates correlations between measured gene expression and assumed sine waves, this method also asks whether gene expression changes according to a given pattern. More specifically, we tested whether the expression profile for each measured transcript across sleep drive states, significantly correlates with a pre-defined template. The template consists of values from 0 to 1, where 0 corresponds to the condition of lowest sleep pressure and 1 to the highest. Another five conditions are assigned to continuous values between 0 and 1 according to the respective level of sleep pressure the animal experienced (**Fig. 5a**, also see Methods). We find that 65.1% of all clusters and 68.4% of annotated clusters have at least one sleep drive correlate (**Extended Data Fig. 9a**).

**Figure 5.**
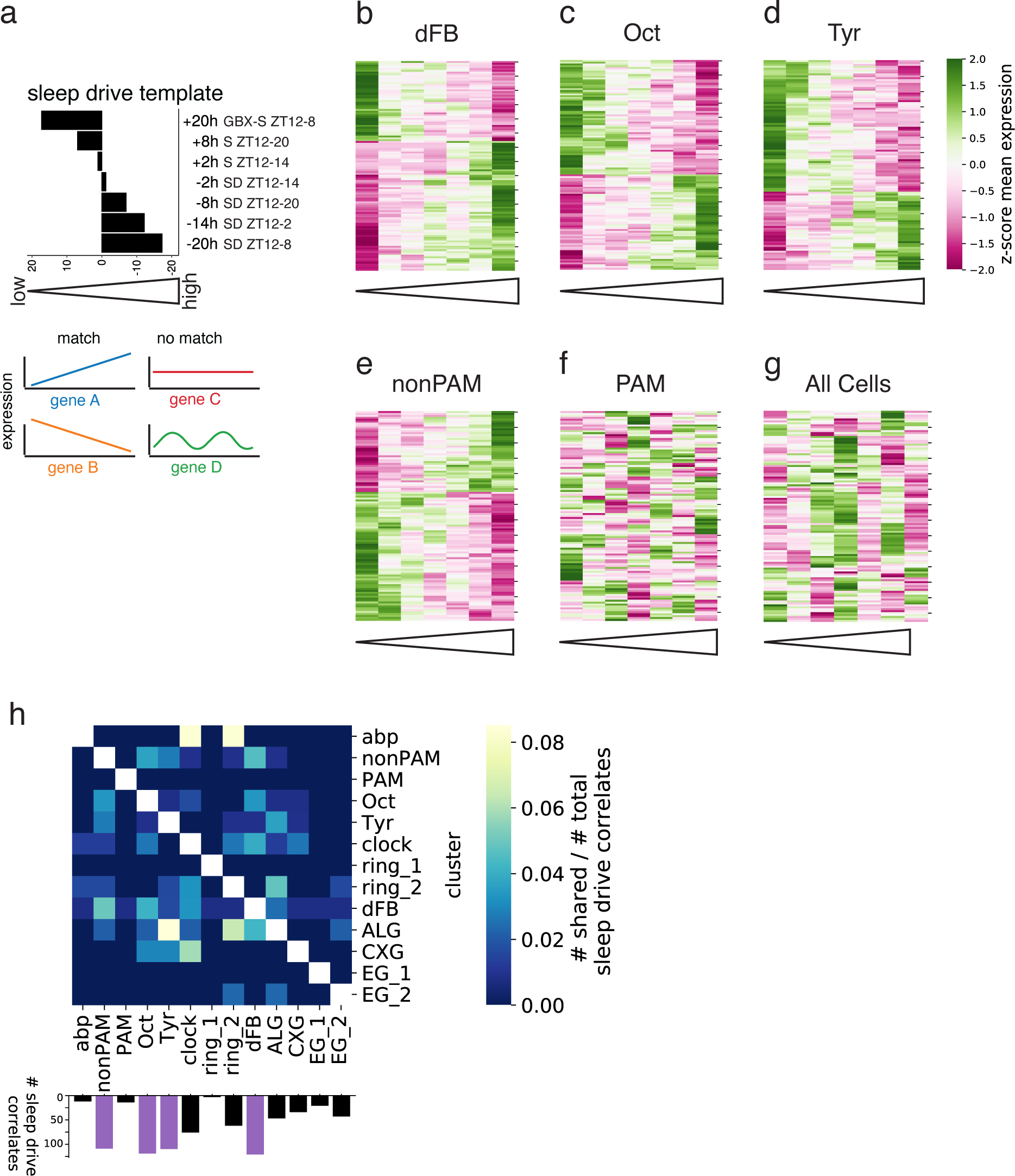
Analysis of molecular correlates of sleep drive. **a.** Illustration of sleep drive template. Each bar represents the amount of sleep or sleep deprivation, that animals experienced prior to sequencing the transcriptome of their central brains. **b-g.** Cluster map of all significant sleep drive correlates of dFB (b), Octopaminergic (c), Tyraminergic (d) and non-PAM dopaminergic neurons (e). Gene expression of sleep drive correlates from non-PAM dopaminergic neurons do not correlate with sleep drive template in PAM dopaminergic neurons or all cells combined (pseudo bulk). **h.** Heatmap of intersecting sleep drive correlates between all annotated cell types with at least one shared gene relative to the total number of sleep drive correlates by cluster.

Interestingly, the four annotated clusters with the highest amount of sleep drive correlates are cell populations known to regulate sleep drive, spearheaded with 121 correlates by the dFB neurons (**Fig. 5h, Extended Data Fig. 9a-b**), that we previously annotated (**Extended Data Fig. 4**). Similarly, the wake-promoting Octopaminergic, Tyraminergic, and non-PAM DAN neurons each have more than 100 sleep drive correlates (**Fig. 5h, Extended Data Fig. 9a-b**). Plotting gene expression of sleep drive correlates from low to high sleep pressure conditions shows clear correlating patterns for dFB, Octopaminergic, Tyraminergic, and non-PAM DAN neurons (**Fig. 5b-e**). In contrast, the related dopaminergic subtype of PAM neurons only has 14 correlates. Furthermore, the expression of non-PAM DAN sleep drive correlates in PAM DAN neurons is not correlating (**Fig. 5f**), as it is not in all cells combined (pseudo bulk) (**Fig. 5g**). Thus, here we demonstrate that the template matching method is sufficiently sensitive and specific to capture sleep drive correlates of previously identified sleep circuits.

Analogous to asking whether circadian cyclers and sleep/wakefulness correlates differ between cell populations, we next assessed whether molecular correlates of sleep drive vary depending on cell identity. We find that the specificity of sleep drive correlates to a cell type is even more pronounced compared to that of the sleep/wakefulness correlates and circadian cyclers. The highest overlap across all clusters is merely 8% between abp-KC and clock or ring_2 and between astrocytes (ALG) and Tyraminergic neurons (**Fig. 5h**). These data suggest that different cells respond to Process S in unique ways.

### The integrator circuit of sleep homeostasis R5 neurons has a high number of sleep drive correlates

We found that one subcluster of EB ring neurons (ring_2) has a substantial number of sleep drive correlates, while the other (ring_1) shows only a few (**Fig. 5h**). Therefore, we asked whether the previously identified sleep drive regulating R5 neurons are part of the ring_2 subcluster. R5 neurons contain only ∼32 cells in an adult fly brain^26^. To identify them, we first selected all EB ring neurons that were annotated based on previously used marker genes^20^ and repeated dimensionality reduction exclusively on the EB ring neuron cluster to specifically analyze its subclusters. This re-clustering resulted in at least three separated subclusters that express unique transcriptomic signatures (**Fig. 6a**). To identify R5 neurons among the three clusters, we focused on a subset of marker genes (**Fig. 6b**), whose expression pattern is known from T2A-Gal4 knockin driver lines^42, 43^. By crossing relevant driver lines to chemically tagged effectors, we were able to visualize the expression pattern of multiple marker genes in the EB ring neuron subclusters (**Fig. 6c**). EB ring neuron subtypes can be distinguished by their projection patterns into the ring-shaped EB structure^45^. This allowed us to map the subtype identities to some of our single cell EB ring neuron subclusters. Ring_C differentially expresses *cry* and *pdfR* (**Fig. 6b**), two genes that are expressed specifically in neurons projecting into the R5 ring (**Fig. 6c**). Therefore, we identified cluster ring_C as R5. Remarkably, we found a high number of dose-dependent correlating genes specifically in R5 neurons, while few to no genes were identified in the other two subclusters (**Fig. 6d-e**).

**Figure 6.**
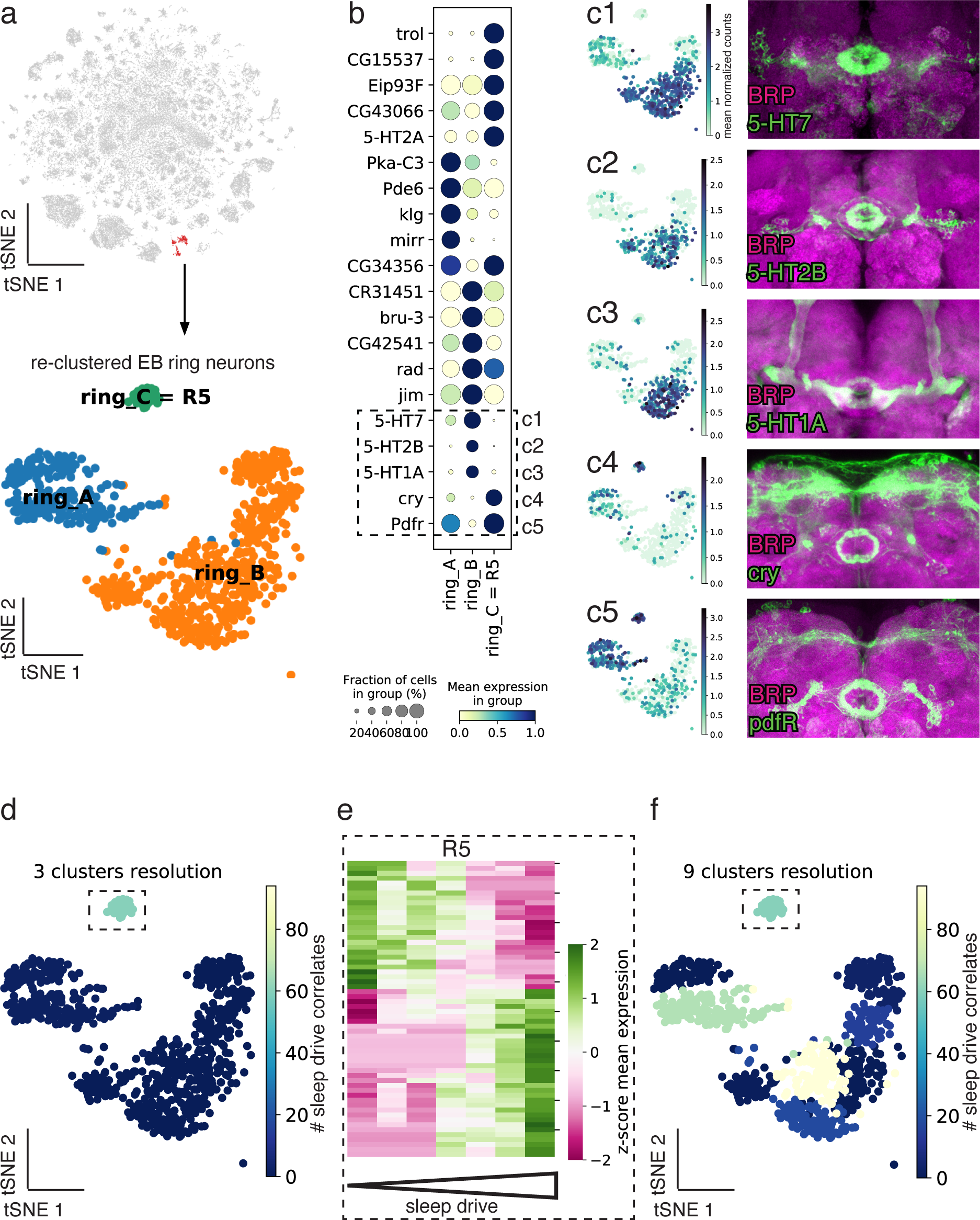
Identification of EB R5 neurons and sleep drive correlates across EB ring neuron subtypes. **a.** Top: tSNE plot of two EB ring neuron subclusters. Bottom: Re-clustering EB ring neurons into three subclusters ring_A, ring_B and ring_C. **b.** Top 20 differentially expressed genes between the three subclusters. **c1-c5.** Expression pattern and chemical tag staining of T2A-Gal4 driver lines of those genes, boxed in (b). Their morphology reveals that ring_C are EB R5 neurons. **d.** tSNE with number of sleep drive correlates at 3 clusters resolution. **e.** Cluster map displaying expression of sleep drive correlates for R5 neurons. **f.** tSNE with number of sleep drive correlates at 9 clusters resolution.

Anatomic analyses have shown that there are eleven subclusters of EB ring neurons^46^. Therefore, we asked whether our two bigger EB ring neuron clusters still contained multiple subpopulations and re-clustered them into another eight subclusters. Matching the sleep drive template to these subclusters revealed that two of them have a high number of correlating genes in addition to R5 (**Fig. 6f**). This is in line with studies that have identified additional ring neuron subtypes apart from R5 that regulate sleep amount and sleep fragmentation^27, 28, 47^. Importantly, the remaining six subclusters had no or few correlating genes with sleep drive (**Fig. 6f**), demonstrating the specificity of the template matching method. This re-clustering analysis also highlights the importance of examining homogeneous populations.

### The homeostatic and circadian processes converge in glial cells

Previously it was shown in flies that the homeostatic and circadian processes interact indirectly by neuronal circuits connecting clock neurons with sleep homeostat circuits (EB ring neurons and dFB neurons)^48–52^. We asked whether there is a more direct interaction of the two processes in the same cell and whether one process affects a cell type more than the other one. To this end, we compared the number of sleep drive correlates with the number of cyclers by cell population and assigned each cell type to either process or to both simultaneously (see Methods). Surprisingly, in most cases, we found that a given cell type was more affected by only one process (**Fig. 7a**). For example, dFB, Oct/Tyr, non-PAM DANs, and R5 neurons have many sleep drive correlates but few circadian cyclers. On the other hand, cell types with many circadian correlates, for example y-KCs, ab-KCs, and PGs, have no sleep drive correlates. Remarkably, two subtypes of EB ring neurons – R5 and ring_B were affected by either process in opposing ways, in accordance with previous findings, showing that the R2/R4m neurons (likely part of ring_B) receive circadian timing information from clock neurons^52, 53^, while R5 neurons themselves encode the sleep homeostat^26^. This suggests that the effect of either the circadian or the homeostatic process differs depending on the cell type and can vary even for closely related cell types. KC subtypes, except for abp-KCs, follow a pattern of high number of circadian “cyclers”, but few or no sleep drive correlates. Intriguingly, both number of cyclers and sleep drive correlates are high in all glia with the exception of PG, as opposed to few neuronal clusters with such high numbers. This demonstrates that a simultaneous convergence of both circadian and homeostatic processes takes place in glial cells, as their transcriptome is affected by both.

**Figure 7.**
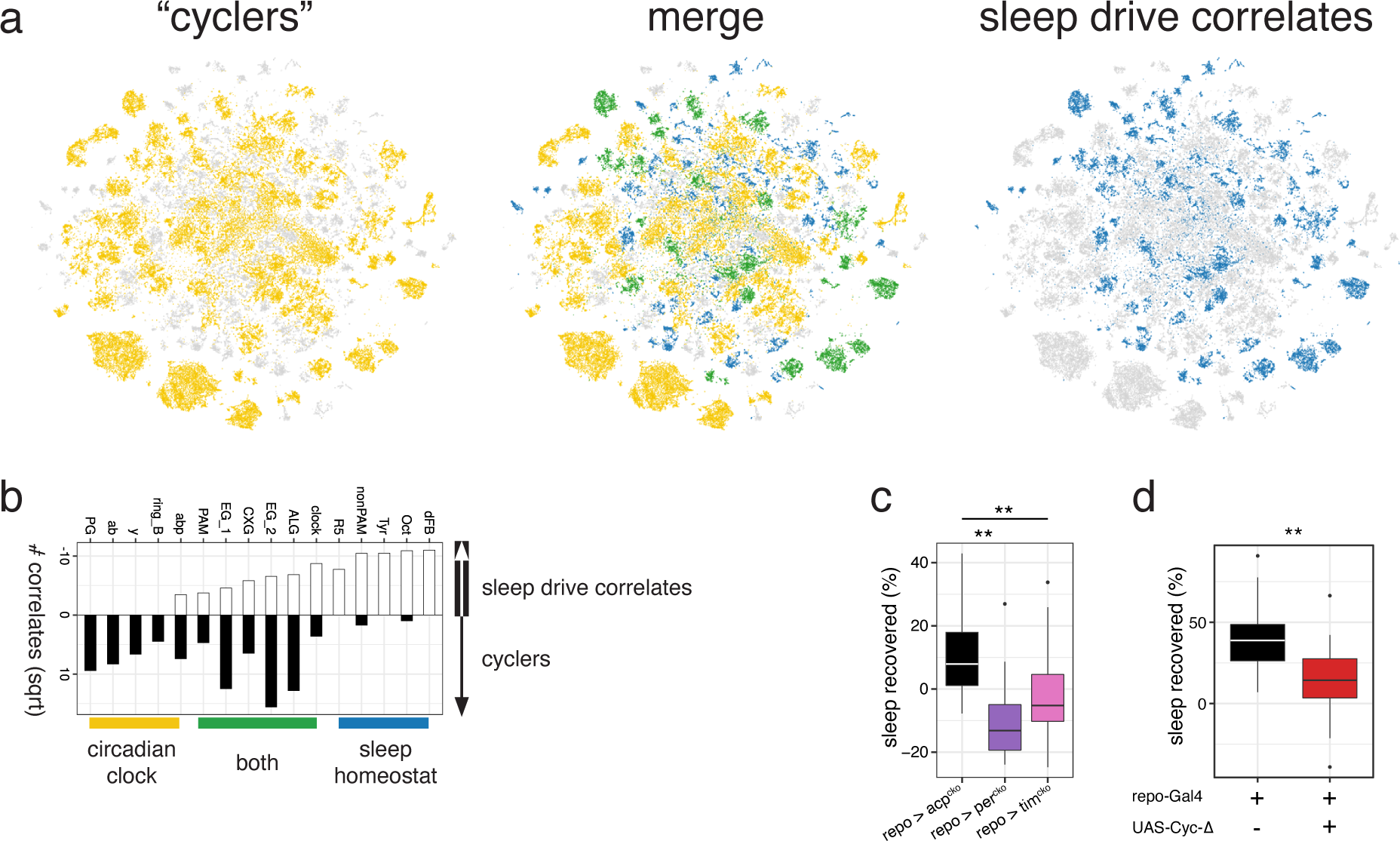
The homeostatic and the circadian process converge on glial cells. **a.** Clusters assigned to “cyclers” (yellow, left) or sleep drive correlates (blue, right) group visualized in tSNE. Middle tSNE shows the merge of left and right tSNE, highlighting the clusters assigned to both groups in green. **b.** Number of correlating genes with circadian or sleep drive template across all annotated clusters that have correlates for either process. Colour indicates their assigned group. **c.** Sleep recovery remains unchanged for 3.5 hours post-SD of lies expressing conditional knock-out of PER (n=10) or TIM (n=22) compared to baseline sleep of the same ly in same ZT time period prior to SD (purple and pink), while control lies show robust rebound sleep post-SD (black) (n=15), p=0.005506 and p=0.007333 (two-sided t-test). The boxplots indicate the minimum, median, maximum, frst quartile and third quartile. Error bars represent the frst (third) quartile - (+) 1.5*IQR (interquartile range). **d.** Sleep recovery remains unchanged for 3.5 hours post-SD of lies expressing dominant-negative forms of eve in glia compared to baseline sleep of the same ly in same ZT time period prior to SD (red) (n=20), while repo-Gal4 > + control lies show robust rebound sleep post-SD (black) (n=15), p=0.002966.

Interestingly, we identified genes that regulate the core molecular clock, *vri* and *CG31324*, as sleep/wakefulness correlates in glia. Similarly, expression levels of *E23*, a regulator of the circadian rhythm, correlate with sleep drive in glia specifically. Thus, Process S may directly influence the core clock machinery in glial cells. Next, we asked whether the disruption of the circadian clock, specifically in glia, would result in impairment of the sleep homeostat.

We expressed a dominant negative form of CYCLE (CYC)^53^ or conditionally knocked out the PER and TIM^54^ specifically in glial cells, while leaving all neurons including clock neurons unaffected. To assess sleep homeostasis in these animals, we sleep deprived flies for 12 hours during the night and measured rebound sleep, a hallmark of sleep homeostasis, in the following morning. Flies with a disrupted glial clock show significantly reduced rebound sleep after sleep deprivation compared to control flies (**Fig. 7c-d**). These data indicate that the glial clock is required for normal sleep homeostasis, and suggest that Process S and Process C directly influence each other in glial cells to determine sleep-wake cycles.

## Discussion

The present study is the first to profile gene expression across different sleep and wakefulness states, degrees of sleep pressure, and diurnal time points in an unbiased manner across all cell populations of an entire central brain. We show that sleep/wakefulness and sleep pressure as well as circadian cycling correlates can be found at the transcriptional level across our 214 clusters, that include 20 annotated clusters. Depending on their cell identity, few correlates overlap between clusters. The specificity of correlates suggests that distinct cell populations are affected differently by sleep/wakefulness, sleep drive, and circadian time. We further show that many correlates are only visible in homogenous populations, not in a bulk sample. This observation likely stems from the finding that correlates are largely unique to a specific (sub)cluster. Particularly for sleep drive correlates, we observed that the higher the granularity of a cluster, for instance the EB ring neurons, the more correlates are captured, highlighting the importance of cluster homogeneity in such analyses. Further support for the notion that molecular correlates are only visible at single cell resolution, and not in a pseudo-bulk sample stems from the observation that the results of previous bulk transcriptomic studies^7, 8^ that have profiled transcriptome changes across the sleep-wake cycle in *Drosophila* have reached non-overlapping results. Similarly, our pseudo bulk sleep-wake correlates only overlap by a maximum of 2% compared to their results (**Extended Data Fig. 10**). Therefore, our data suggest that single-cell transcriptomics enable the discovery of gene expression correlates in different cell types and that these effects are masked in bulk data.

We found that the core clock machinery exists only in clock neurons and glial cells, but not in other neurons in the fly brain. This is consistent with a previous report of the absent expression of core clock genes in *Drosophila* dopaminergic neurons^55^ and Kenyon cells^56^. At the same time, most of the neurons do express cyclers. These cyclers in non-clock neurons are likely driven by cells containing the molecular clock, including clock neurons and glial cells. Glial cells may be a more suitable candidate for this role, considering their large number and coverage across the entire brain in contrast to a mere ∼150 clock neurons, whose processes cover only a small proportion of the brain. Supporting this idea, the glial clock in mice has been shown to drive the circadian clock gene expression in the SCN and is sufficient to initiate and sustain circadian locomotor rhythms^57^. Similarly, in flies perturbing glial release results in loss of circadian rhythmic morphological plasticity of pacemaker small ventral lateral neurons^34^. Thus, the circadian cyclers in many non-clock neurons in flies may be driven by the molecular clock in the nearby glial cells.

By using single-cell transcriptomics and template-matching, we detected gene expression changes associated with different sleep drive levels in real-time while sleep pressure accumulates for all cell types separately. Surprisingly, many clusters show a high number of transcriptional sleep drive correlates, including known sleep homeostasis regulating circuits, such as R5 and dFB neurons which are directly involved in sleep homeostasis, as well as the wake-promoting Tyraminergic, Octopaminergic and non-PAM dopaminergic neurons that interact with the sleep homeostat. Furthermore, many sleep drive correlates are captured in glial cells, several subtypes of which have previously been shown to regulate sleep/wakefulness or sleep homeostasis in *Drosophila*^25, 58^. Considering the specificity of the method to identify relevant sleep homeostasis regulating circuits, other unannotated clusters with a high number of sleep drive correlates may also be involved in the homeostatic regulation of sleep, and it will be interesting to determine their identity to ultimately examine their sleep-regulating function.

Sleep homeostasis and circadian rhythms are two distinct behavioural processes and likely affect different cells in the brain. In accordance, we found that most cells in the fly brain have either a high number of sleep drive correlates but a low number of circadian cyclers, or the reverse. However, while the homeostatic and circadian processes are known to function independently, increasing evidence suggests a cross-talk between these two processes^59, 60^. In flies, recent studies suggest a model where the sleep homeostat and the circadian clock interact indirectly with each other through neuronal circuit connections. Several circuits have been shown to convey the circadian time information from clock neurons to the sleep homeostat centres EB and dFB^48, 49, 53^. Reciprocally, hugin+ neurons act downstream of dFB neurons and modulate clock neurons^50^. However, our data argue for an alternative model, in which the sleep homeostatic and circadian process interact directly in glial cells to regulate the sleep-wake cycle. In this model, the sleep-wake cycle affects the regulators of core clock genes specifically in glia, but not in clock neurons. At the same time, the molecular clock in glia is required for sleep homeostasis. How do these two processes interact in glial cells? We and others previously demonstrated that glial Ca^2+^ signalling encodes the level of sleep need^25, 58^. In addition, Ca^2+^ signalling plays an important role in regulating the oscillation of core clock genes and many Ca^2+^ channels and transporters are rhythmically expressed in mammalian clock neurons^61^. Thus, the reciprocal interaction between Ca^2+^ signalling and the molecular clock in glia may be the molecular substrate of the interaction of homeostatic and circadian process to ultimately instruct downstream neurons and appropriate behaviour.

## Methods Animals

The following *Drosophila* strains were ordered from the Bloomington *Drosophila* Stock Center (BDSC): DGRP (*Drosophila* Genetic Reference Panel) 88, 287, 303, 313, 359, 379, 441, 646, 892 and 908; 5-HT1A-Gal4 (#84588); Pdfr-Gal4 (#84684); cry-Gal4 (#24514); UAS-Cyc Δ (#36317), iso31 (#5905), repo-Gal4 (#7415), UAS-sgRNA-tim3x (#90768), UAS-sgRNA-per4x (#90769), UAS-sgRNA-acp98AB4x (#90770). 5-HT2B-Gal4 and 5-HT7-Gal4 were a gift from Hiromu Tanimoto.

### Sleep behaviour

After entraining male flies for three days, four to nine days-old single flies were loaded during the active period between ZT 0-2, and sleep was recorded in 12:12 LD condition in a constant environment of 55-65% humidity and 22℃. Locomotion was tracked in a high-resolution video-based Raspberry Pi-enabled device (ethoscope)^62^. Targeted mechanical sleep deprivation was delivered in a feed-back loop triggered by 10 seconds of quiescence of a fly by the rotation of the respective tube. Sleep analysis was performed with adaptations to the rethomics pipeline in R.

For sleep profiling DGRP lines, thirty-six DGRP lines were selected based on sleep architecture metrics previously reported in 168 DGRP lines^17^. Of the thirty-six lines we thoroughly compared sleep amount, latency, fragmentation, and depth. We selected those lines with sleep patterns similar within and between the lines. Principal Component Analysis and analysis of pDoze and pWake^18^ were performed in R. Plots in **Extended Data Fig. 2** were made in Python and R. Those flies that slept less than 50% of the average amount of their genotype were excluded (4.5% - 37/812 flies).

For single cell sequencing experiments, around 200 flies were loaded belonging to different combinations of the behavioural conditions linked to one or two of the ten DGRP lines in each of seven runs. Flies belonged to two to four conditions and five to nine DGRP lines. The link between condition and DGRP line was shuffled in every run. Flies were pre-selected based on the recorded baseline sleep and wake behaviour on the day prior to dissection.

Immediately prior to dissection, a final selection of 40 flies was decided based on the previous day’s pre-selection and the sleep or wake behaviour up to the point of dissection in real-time. Dissection, dissociation and 10x processing followed immediately after selection.

Sleep rebound was assessed after 12h targeted sleep deprivation between ZT12-ZT24 and two prior baseline nights in 4-7 days old mated females. Rebound was calculated as the difference between the mean fraction asleep during the first 3.5 hours post-SD from ZT0-ZT3.5 and the fraction asleep during the same ZT period post second baseline night. We excluded animals that slept more than 0.2% of the sleep deprivation period and if they slept less than 50% of the amount of average night-time sleep of their genotype.

### Brain dissociation into single cells

In each of seven replicates, around forty central brains were dissected in ice-cold Schneider’s Medium with 30 μM of the transcription inhibitor Actinomycin D (Sigma-Aldrich 1410).

Within 45 min after starting dissections, the brains were dissociated with an enzyme mixture of dispase (3mg/mL), collagenase (100mg/mL) and trypsin-EDTA (0.05%) in a Thermoshaker at 25C for 15 min. The dissociation was reinforced by pipetting the solution at least four times during the incubation. After washing with DPBS 15 μM Actinomycin D, resuspending the cell pellet in 100-200 ml DPBS 0.04% BSA and filtering the cell pellet with a 10 μM pluriStrainer (ImTec Diagnostics_435001050).

### 10x Genomics

Library preparations for the single cell RNA-seq was performed using 10X Genomics Chromium Single Cell 3’ Kit, v3.1 NextGEM chemistry (10X Genomics, Pleasanton, CA, USA). The cell count and the viability of the samples were assessed using LUNA dual florescence cell counter (Logos Biosystems). For each sample a targeted cell recovery of 10000 cells was aimed for. The dissociated cells from a fly brain are typically quite small (averaging between 1-4 um) and we have seen that this affects the counting accuracy. In order to get consistent counting accuracy, the average of cell counts from three independent measurements were consider for loading the fly cells to the 10X controller. Post cell count and QC, the samples were immediately loaded onto the Chromium Controller. Single cell RNAseq libraries were prepared using manufacturers recommendations (Single cell 3’ reagent kits v3.1 user guide; CG000204 Rev D), and at the different check points the library quality was accessed using Qubit (ThermoFisher) and Bioanalyzer (Agilent). For a targeted sequencing saturation of 50-60%, sequencing was performed at a depth of 30,000 - 60,000 reads per cell and single cell libraries were sequenced either on Illumina’s NovaSeq 6000 platform or HiSeq 2500 platform using paired-end sequencing workflow and with recommended 10X; v3.1 read parameters (28-8-0-91 cycles).

### 10x Genomics Data Pre-Processing

The 10x data was mapped to the Drosophila Melanogaster (BDGP6 assembly) genome using CellRanger v.3.1.0. SNP data for the DGRP lines was downloaded from http://dgrp2.gnets.ncsu.edu/ and liftOver was used to convert these to dm6 coordinates, SNPs were then filtered to include only those present uniquely in one of the lines used. This vcf file was used to run demuxlet from the popscle package [https://github.com/statgen/popscle] with default parameters, tools from https://github.com/aertslab/popscle_helper_tools were utilized to speed up computation.

## Data Processing

Scanpy (v1.4.4) was used to process the 10x libraries, samples were first loaded and cells were annotated with genotypes determined by demuxlet. Demuxlet determined doublets and ambiguous cells were removed, any cells assigned to a genotype not present in that experiment were also removed. Finally, scrublet was used to remove remaining doublets using an expected doublet rate determined by the following formula based on the known doublet rate of the 10x chromium device and the number of doublets expected to be remaining after demuxlet (within-genotype doublets): (0.008 * (n_cells / 1000)) * (num_lines / (num_lines ^ 2)).

Following doublet removal, cells with less than 200 genes were removed, as well as cells with more than 20-30% UMIs assigned to mitochondrial genes (thresholds can be found in **Supplementary Table 4**), cells belonging to the ZT2 Wake condition were also removed. All remaining cells were combined into a single sample for further processing. Cells were normalized to a total of 10000 counts per cell and log transformed, highly variable genes were identified using default parameters and number of counts and percentage mitochondrial reads were regressed out, finally, counts were scaled to unit variance with a zero mean with a max value of 10. A principal component analysis was performed on the data and the pcacv workflow from vsn-pipelines (v0.25.0) [https://doi.org/10.5281/zenodo.3703108] was used to determine the number of principal components (PCs) (58 PCs) to continue with for further dimensionality reduction and clustering. Both UMAP and tSNE dimensionality reductions were computed and clusters were determined using the louvain algorithm at various resolutions (**Supplementary Table 2**). Final anndata objects were converted to loom files for visualization in SCope (https://scope.aertslab.org).

### Multiplexing strategy

DGRP lines can be distinguished from one another by their unique SNPs. Their natural genetic variation from each other allows us to determine the condition for each cell from the sequenced data^13–15^. This strategy has the added advantage of removing around 90% of droplets that enclose two cells instead of one from the dataset. This multiplexing strategy minimizes technical variation and batch effects, otherwise arising from separate cDNA library preparations.

### Integration of dorsal fan-shaped body neurons (dFB) SMART-seq2 data

To identify the cell cluster containing dFB neurons in our dataset, we made use of the publicly available transcriptome data of a FAC-sorted dFB subset that was genetically targeted with the R23E10-Gal4 driver. The raw data that was processed with a standard SMART-seq2 protocol was downloaded from GEO (Accession: GSE107451)^19^. Reads were cleaned using fastp (v0.20.0) [PMID: 30423086] and mapped to the BDGP6 Drosophila Melanogaster and quantified using STAR (v2.7.9a) [PMID: 23104886]. A non-negative least squares (NNLS) regression model was used to match dFB SMART-seq2 cells to their corresponding cluster in our atlas (**Extended Data Fig. 4**), and was performed as previously described^19^, with the top 10 marker genes (determined by z-score) per cluster from resolution 0.8 being used as the gene subset.

### Downstream analyses

All analyses (differential expression analysis, sleep drive template matching and identification of cycling genes) except for the EBM classifier were performed on all clusters in high resolution 8.0 of the louvain clustering, except for those clusters annotated in a different resolution (**Supplementary Table 2**). Also, we excluded genes that displayed obvious expression differences between different runs for all analyses (**Supplementary Table 3**).

### Validation of circadian clock related genes with SCENIC

Gene co-expression analysis was performed with SCENIC (single-cell regulatory network inference) using the scenic_multiruns workflow from vsn-pipelines in two steps (to decrease computational time). First, 10 runs were performed with a list of all transcription factors, any transcription factors linked to a motif or track that were detected as a regulon by SCENIC were used to perform a second step of an additional 90 SCENIC runs only looking for these transcription factors. The AUC value quantifies the presence of that motif in a cell type. We compared AUC values of the *Clk* regulon across all cell types.

### Identification of cycling genes

To identify cycling genes between ZT 2, 8, 14 and 20, we used the JTK algorithm of the MetaCycle package^63^. As MetaCycle takes a matrix of three replicates for each time point, we treated each cell in a cluster as a replicate and randomly assigned it to one of three pseudo-replicates for each ZT timepoint. The mean expression value for each combination of cluster and ZT timepoint replicate was calculated by transcript. This matrix served as input for the JTK algorithm. The matrix was filtered to test only genes that had a maximum expression of 0.8 CPM and an expression amplitude of at least 1.5-fold between time points, similar to a previous study^29^. We repeated the random allocation three times. Only clusters for which the matrix generation was successful in all three technical replicates, were considered for further analysis, leaving 183 of the 214 initial clusters. The matrix generation was successful in all annotated clusters. A gene was labelled as significantly cycling (Benjamini-Hochberg corrected two-tailed p-value of 0.05) in a cluster only if it was detected as such in all three technical replicates. The threshold set to assign a cluster into the “cyclers” group (**Fig. 7a**) was 3, which is the mean of the square root of the number of cycling correlates for all 195 clusters.

### Differential Expression Analysis

The Wilcoxon rank sum test was used to compare groups of cells for differential expression analysis, this was performed using the rank_genes_groups function from Scanpy where n_genes equalled the number of genes in the dataset, reference and test groups were set per DE analysis. Transcripts were considered differentially expressed genes (DEG) if their Benjamini-Hochberg FDR corrected p-value was less than 0.05.

All cells except for KCs were divided into either cholinergic, GABAergic, glutamatergic or unknown category based on their expression of *VAChT*, *Gad1* and *VGlut* above a value of 0.5, 0.4 and 0.5, respectively.

### Tree-based modelling of sleep and wake states

We performed a binary classification task to learn to map the transcriptome of single cells to the behavioural labels “wakefulness” or “sleep”. Specifically, we trained separate instances of the Explainable Boosting Machine, using identical training settings and hyperparameters, implemented in the interpret.glassbox.ebm.ExplainableBoostingClassifier class with default settings as or something declared in scikit-learn 1.0.2 and interpret 0.2.7. Each instance was trained on a different cell subtype within a group of related cell types or “backgrounds”, specifically the subtypes of glia or KCs. Marker genes were excluded from the training to prevent classification based on cell identity as opposed to behavioural state. Marker genes were determined by performing differential expression analysis between either the glia or KC subtypes. This analysis resulted in excluding those genes with a log fold change larger than 3.5 (glia) and 2 (KC) during training (**Extended Data Fig. 7a-b**). The trained models were then used to predict behavioural state on test datasets of each glial or KC subtype to compare classification performance on same and different cell subtypes.

### Finding shared genes between DEA and EBM

The cut-off of top features that resulted from the EBM classifier outlined above was decided by calculating the mean + 2*standard deviation across all features per subtype. Features falling above this threshold were regarded as significantly contributing to the classifier’s decision. The same thresholding was applied to the features used by the control classifier that predicted cell subtype identity. The resulting control features were then merged with the feature subsets generated in the first step. The remaining genes were then merged with all significant corrected p-values of DEGs from the differential expression analysis (see above) on either glia or KCs.

### Sleep drive template matching

Expression of all transcripts across different conditions were tested for a significant Pearson r2 correlation with a sleep drive template for each cluster separately. Sleep drive correlates below a Benjamini-Hochberg FDR corrected p-value of 0.05 were considered significant.

The template consists of 7 conditions ordered by their respective amount of sleep or sleep deprivation. A value between 0 and 1 was assigned to each condition according to that order with even intervals (see **Fig. 5**). The gene expression matrix generation across the 7 conditions failed for 19 of the 214 clusters, including none of the annotated clusters. For the remaining 195 cell populations, for each transcript it was assessed whether there is a significant correlation between the sleep drive template and the expression value of each gene across the sleep and sleep deprivation conditions (**Fig. 5a**). The threshold set to assign a cluster into the sleep drive correlates group (**Fig. 7a**) was 3.72, which is the mean of the square root of the number of sleep drive correlates for all 195 clusters.

### Single-molecule fluorescent in-situ hybridisation

Custom probes with hybridization chain reaction (HCR) technology were acquired from Molecular Instruments, Inc. The protocol was optimized based on the protocol for zebrafish larvae from Molecular Instruments, Inc. and a whole-mount adult fly FISH protocol^64^. Briefly, brains were dissected in S2 medium, fixed in 2% PFA for 55 min, washed with 0.5% PBST 3 x 15 min, sequentially dehydrated and incubated overnight in 100% EtOH. After rehydration, brains were washed with 5% SSC 0.1% TritonX 4 x 15 min at RT. Subsequently, they were incubated in probe hybridization buffer at 37 C on a rotator for 1.5h and incubated in probe hybridization buffer and relevant probes overnight. Probes were washed with probe wash buffer 5 x 15 min and 5 x 15 min with 0.1% 5xSSC at RT. Then, brains were incubated for 30 min at RT in amplification buffer and overnight in heat-shocked amplification probes diluted in the same buffer at RT on a rotator. After incubation, brains were washed 5 x 15 min in 0.1% 5xSSC and finally incubated in Vectashield at 4 C overnight before mounting and imaging with a Nikon NiE A1R confocal microscope.

### Whole-mount brain staining

After dissection in S2 medium, fly brains were fixed in 4% paraformaldehyde for 30 min. Then, brains were washed in 0.5% PBST twice for 15 min. Subsequently, brains were incubated in SNAP tag at a dilution of 1:1000 in 0.3% PBST for 30 min and a further 30 min with the addition of the Halo tag at the same dilution. Lastly, brains were washed in 0.5% PBST twice for 15 min and incubated in Vectashield overnight at 4C. Brains were mounted in Vectashield and imaged on a Zeiss Airyscan 880 confocal microscope.

## Data and code availability

The scRNA-seq data and code will be stored online and available upon publication. Gene expression across all cell populations are visualized at https://scope.aertslab.org/#/Fly_Brain_Sleep/Fly_Brain_Sleep%2FFly_Sleep.loom/gene. Transcript expression plots for all correlates and clusters are available at https://joana-dopp.shinyapps.io/Fly_Sleep_Single_Cell/. Both links can be found at https://www.flysleeplab.com/scsleepbrain.

## Acknowledgements

We thank Stein Aerts and members of the SL lab for discussions and specially Miranda Dyson and Lisa van Ninhuys for help with fly brain dissections. Imaging and RNA sequencing were supported by the light microscopy and nucleomics expertise units at the VIB-KU Leuven Center for Brain & Disease Research. Computing was performed at the Vlaams Supercomputer Center (VSC). Fluorescent in-situ hybridization experiments were made available through VIB Tech Watch Funding. We also thank the Bloomington Stock Center and Shu Kondo and Hiromu Tanimoto for providing fly stocks used in this study. This work was funded by a starting grant of the European Research Council (#758580). J.D. holds a PhD Fellowship of the Research Foundation - Flanders (FWO, 11D8820N and 11D8822N). We thank Ibrahim Taskiran for discussions on applying machine learning to our data. We are thankful for helpful comments on the manuscript from Stein Aerts, Mark Wu, Patrik Verstreken, Chien-Chun Chen, Natalie Kaempf and Nicola Fattorelli.

## Author Contributions

J.D and S.L. conceived the study; J.D., E.S.B. and A.O. established sleep behaviour measurements and analysis with ethoscopes; J.D. performed behaviour, imaging and validation experiments; S.K.P. performed 10x genomics; J.D., K.D. and A.O. analyzed scRNAseq data; J.D. and S.L. wrote and revised the manuscript.

---

**Extended Data Figure 1.**
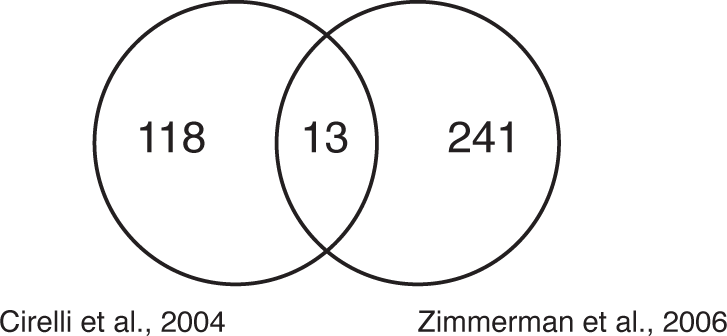
Results of previous bulk microarray studies do mostly not overlap. Transcripts identified in two previous bulk microarray studies that have profiled sleep and wake states of adult fly brains are largely non-overlapping.

**Extended Data Figure 2.**
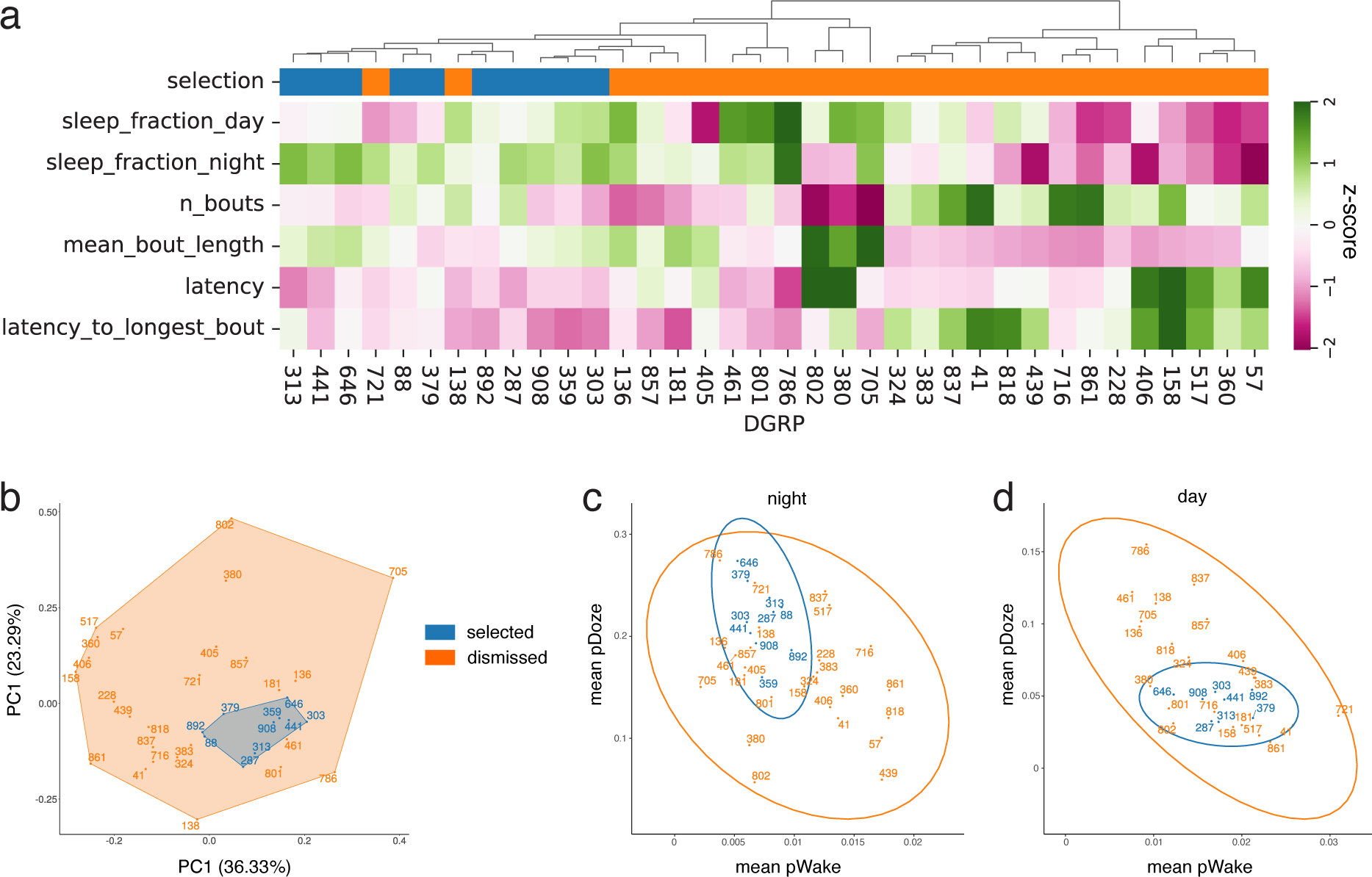
Sleep behavior screening of 36 DGRP lines to select suitable DGRP lines for single cell transcriptomics. **a.** Clustered heatmap of quantified sleep amount during day and night, sleep bout length and number, latency to first and longest sleep bout during nights across all tested DGRP lines. Based on these parameters ten DGRP lines were selected (n=355, yellow) and 26 dismissed (n=425, purple). **b.** Principal Component Analysis on mean and standard deviation per sleep parameter and DGRP line. **c-d.** Scatterplot showing the probability to switch from a wake to a sleep state (pDoze) or vice versa (pWake) averaged across flies of the same DGRP line.

**Extended Data Figure 3.**
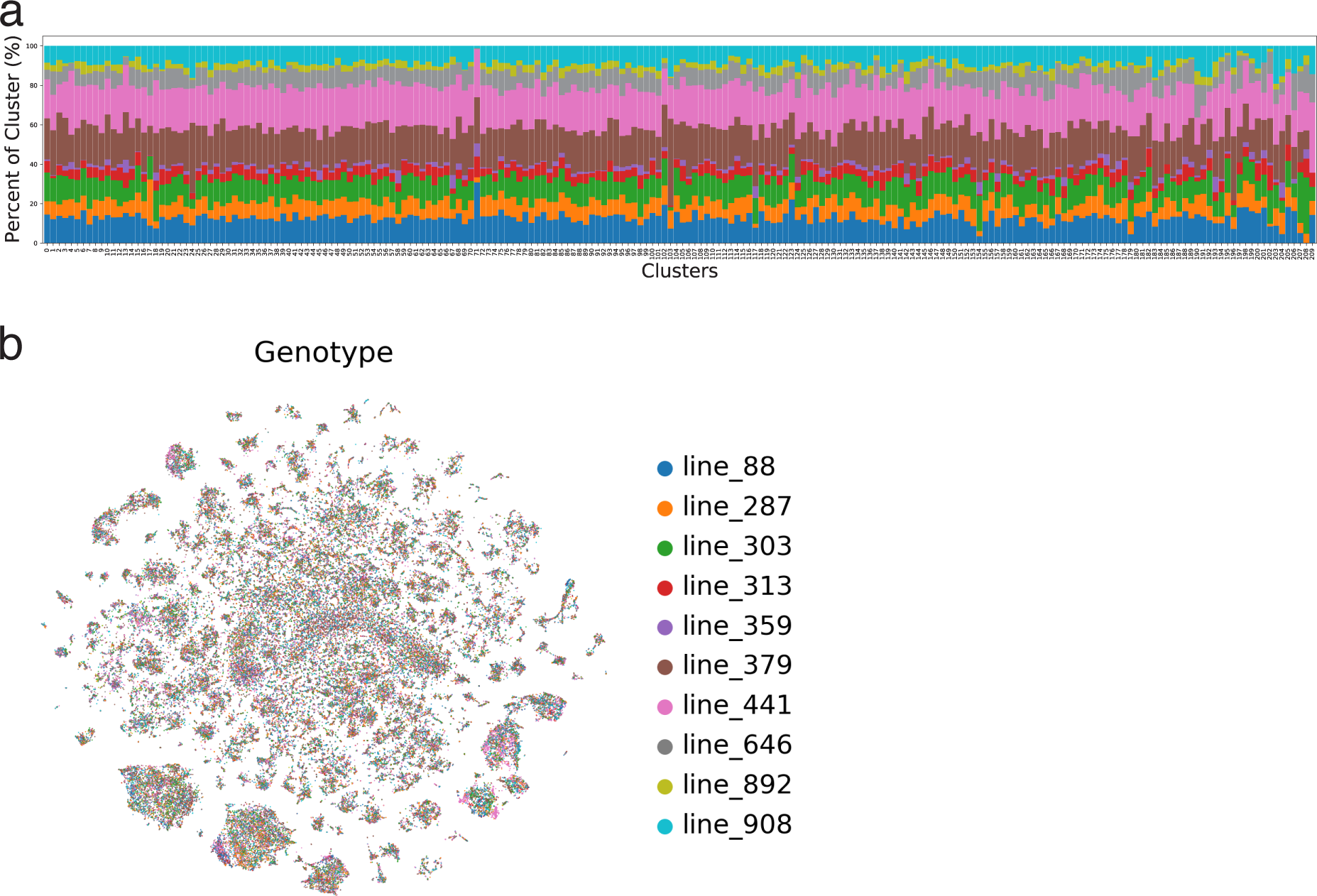
Bias analysis of clusters. **a.** Normalized number of cells coloured by genoype for each cluster in resolution 8. Most clusters have a relatively similar number of cells from each genotype. **b.** tSNE of all cells coloured by genotype shows cells of different DGRP lines mix well.

**Extended Data Figure 4.**
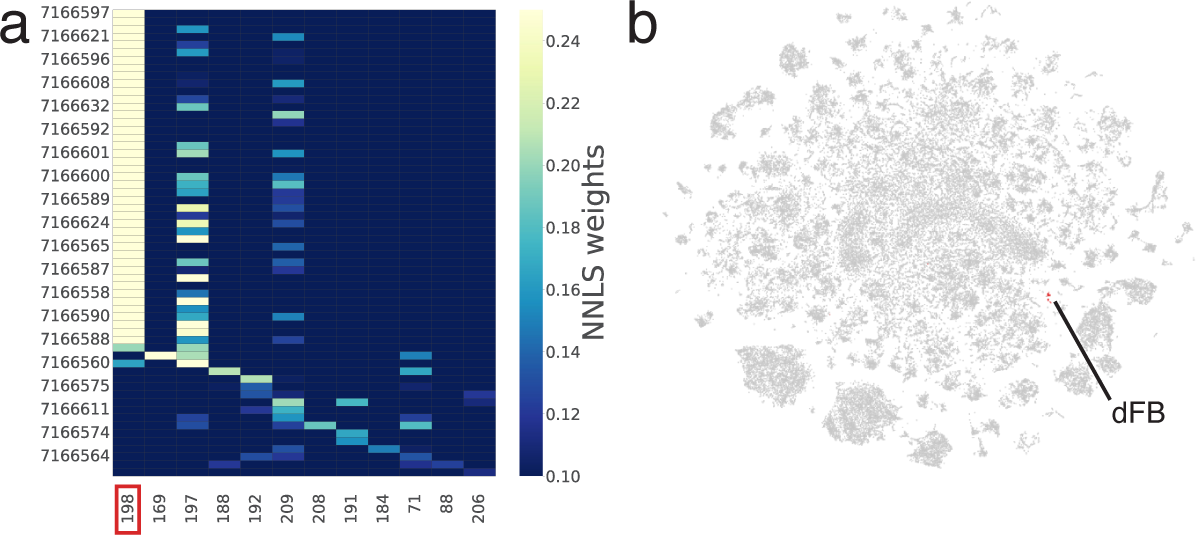
Annotation of dFB cluster. **a.** The 12 clusters of our 10x data, showing a high NNLS score indicating a match with R23E10-Gal4 FAC-sorted scRNA-seq data. **b.** Annotation of cluster 198 as dFB in tSNE of all cell types.

**Extended Data Figure 5.**
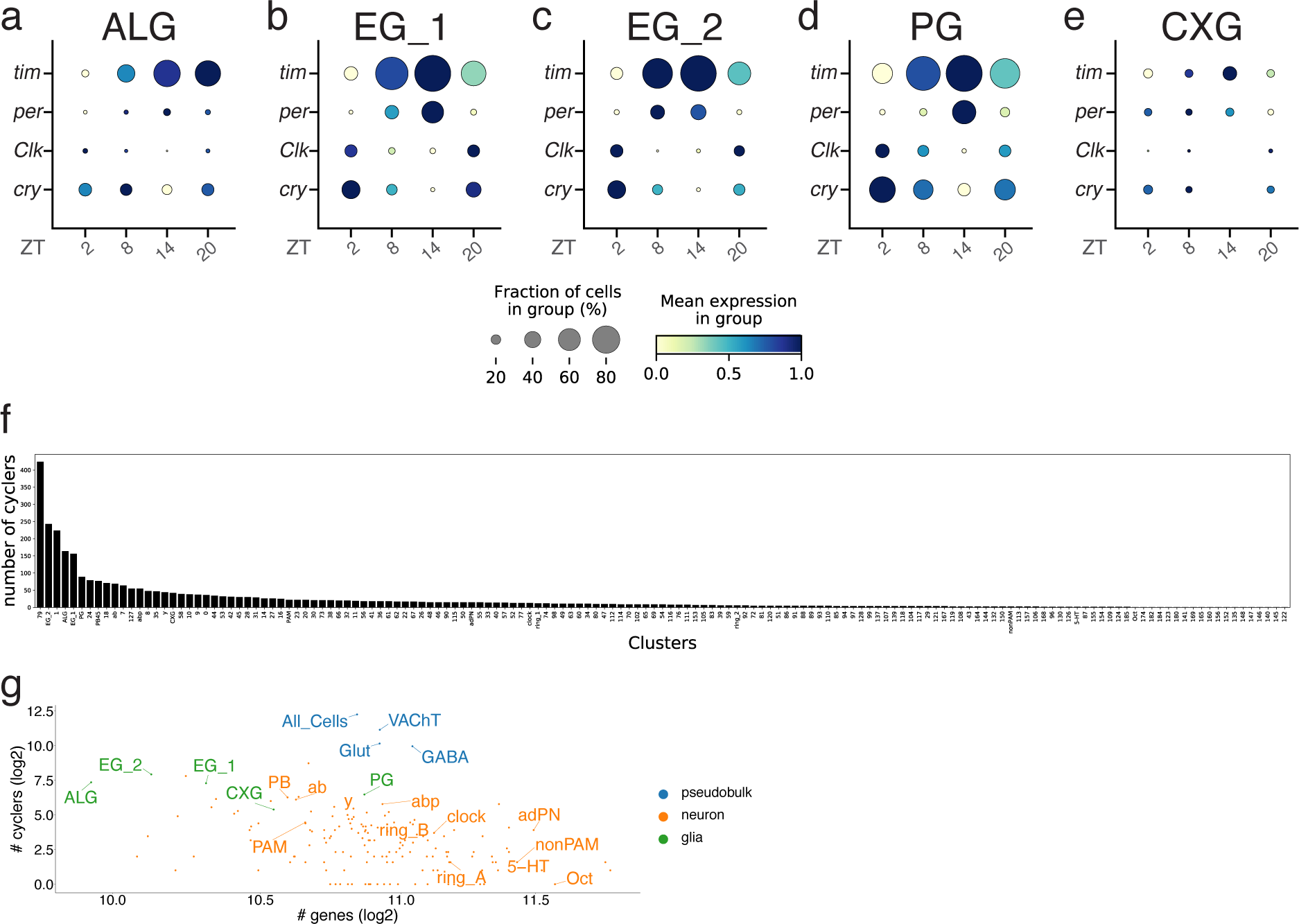
Oscillation of core circadian genes between ZT times in glial subtypes and cyclers across all clusters. **a-e.** Circadian expression levels of core clock genes per, tim, cry and Clk for each glial subtype. **f.** Number of cyclers across all clusters. **g.** Correlation between number of genes and number of cyclers for pseudobulk samples, neuronal and glial cell types.

**Extended Data Figure 6.**
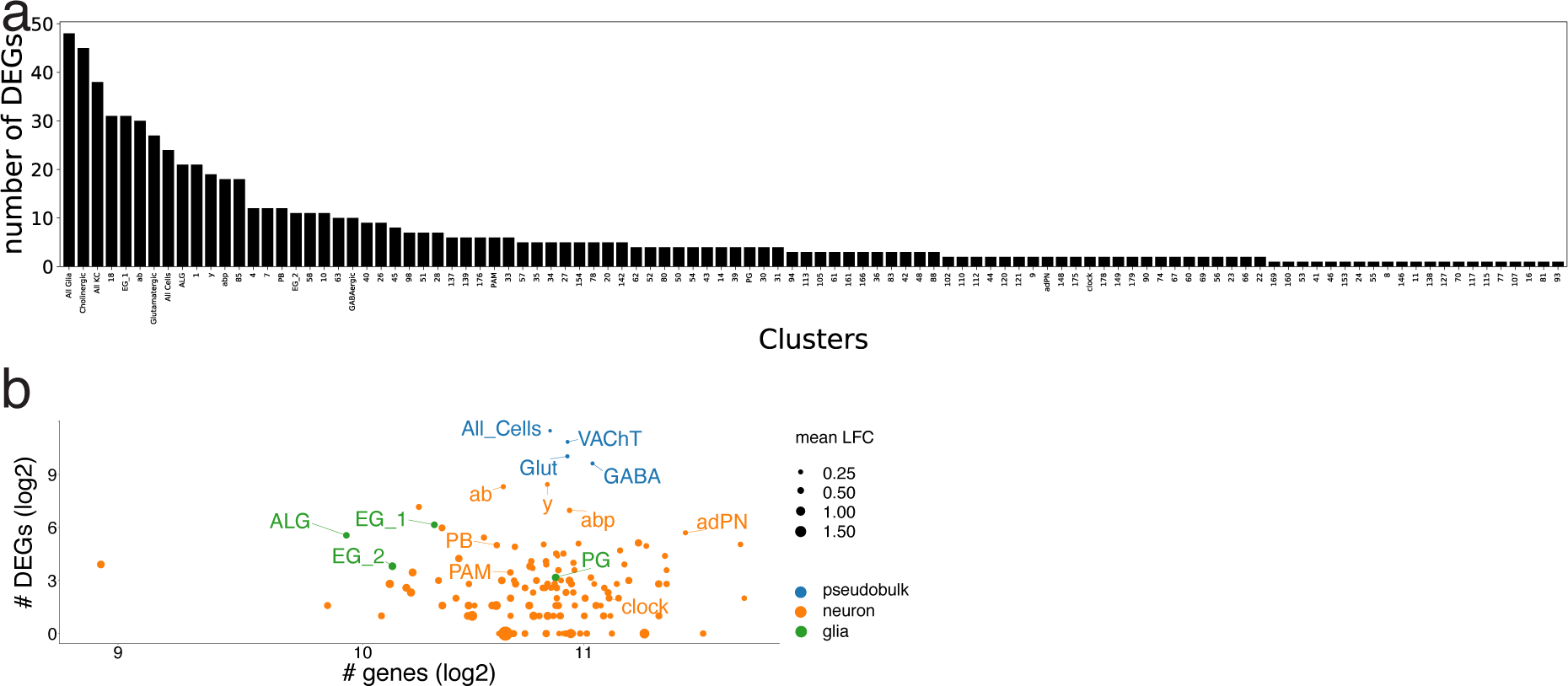
Sleep/wakefulness correlates across all clusters. **a.** Number of DEGs across all clusters. **b.** Correlation between number of genes and number of sleep/wakefulness correlates for pseudobulk samples, neuronal and glial cell types. Dot size indicates average log2 fold change (LFC) across all significant DEGs for the respective cluster.

**Extended Data Figure 7.**
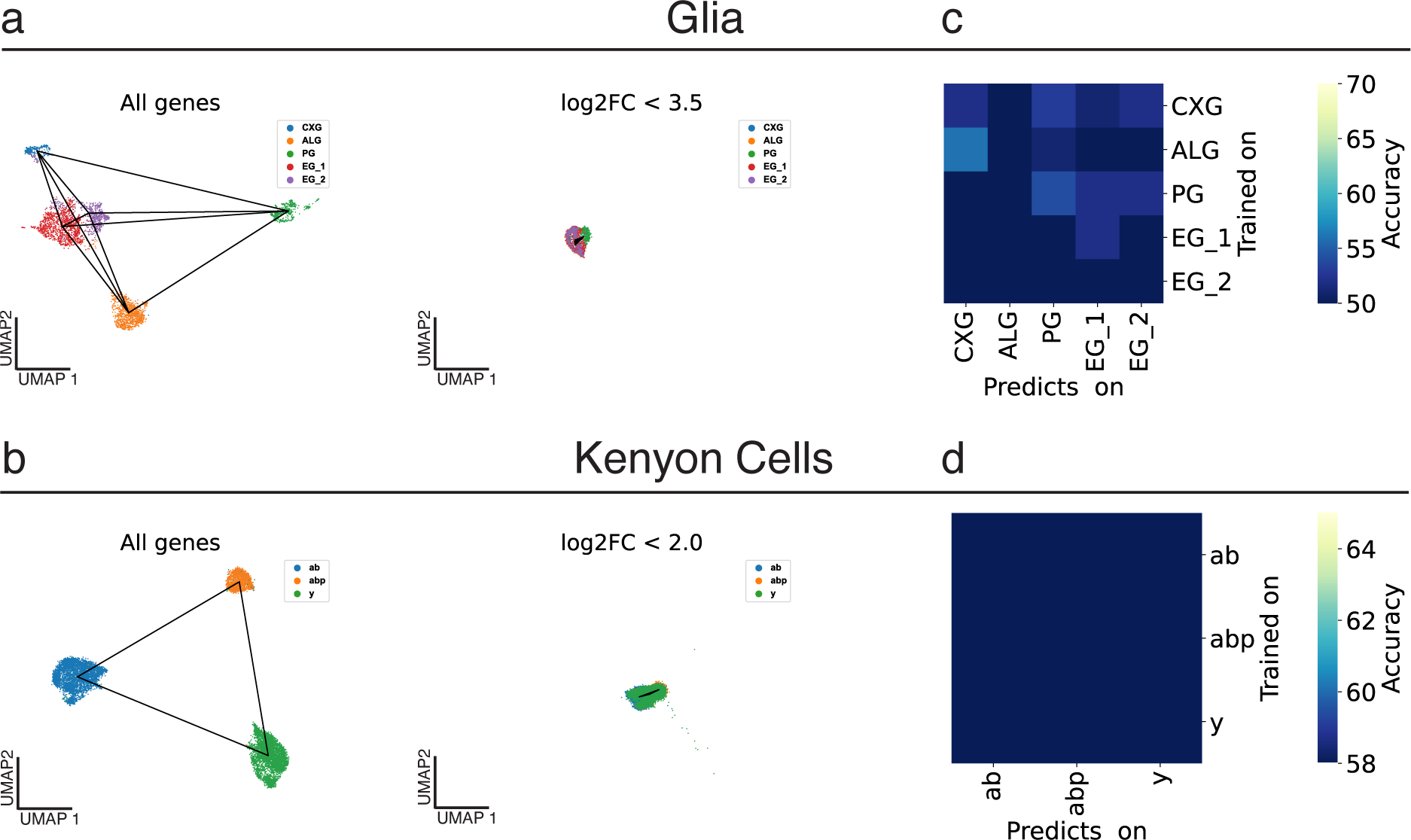
Controlling sleep/wake state classifier by removing marker genes between subtypes of KC or glia. **a-b.** Glial and KC subtypes move closer to each other in a UMAP space, because marker genes above certain log foldchange thresholds are removed. Genes falling below a threshold of 3.5 or 2 for glia and KC, respectively were excluded for training the tree-based EBM classifier. **c-d.** Assigning the sleep or wakefulness label randomly results in random performance of the classifier for the same cell subtype.

**Extended Data Figure 8.**
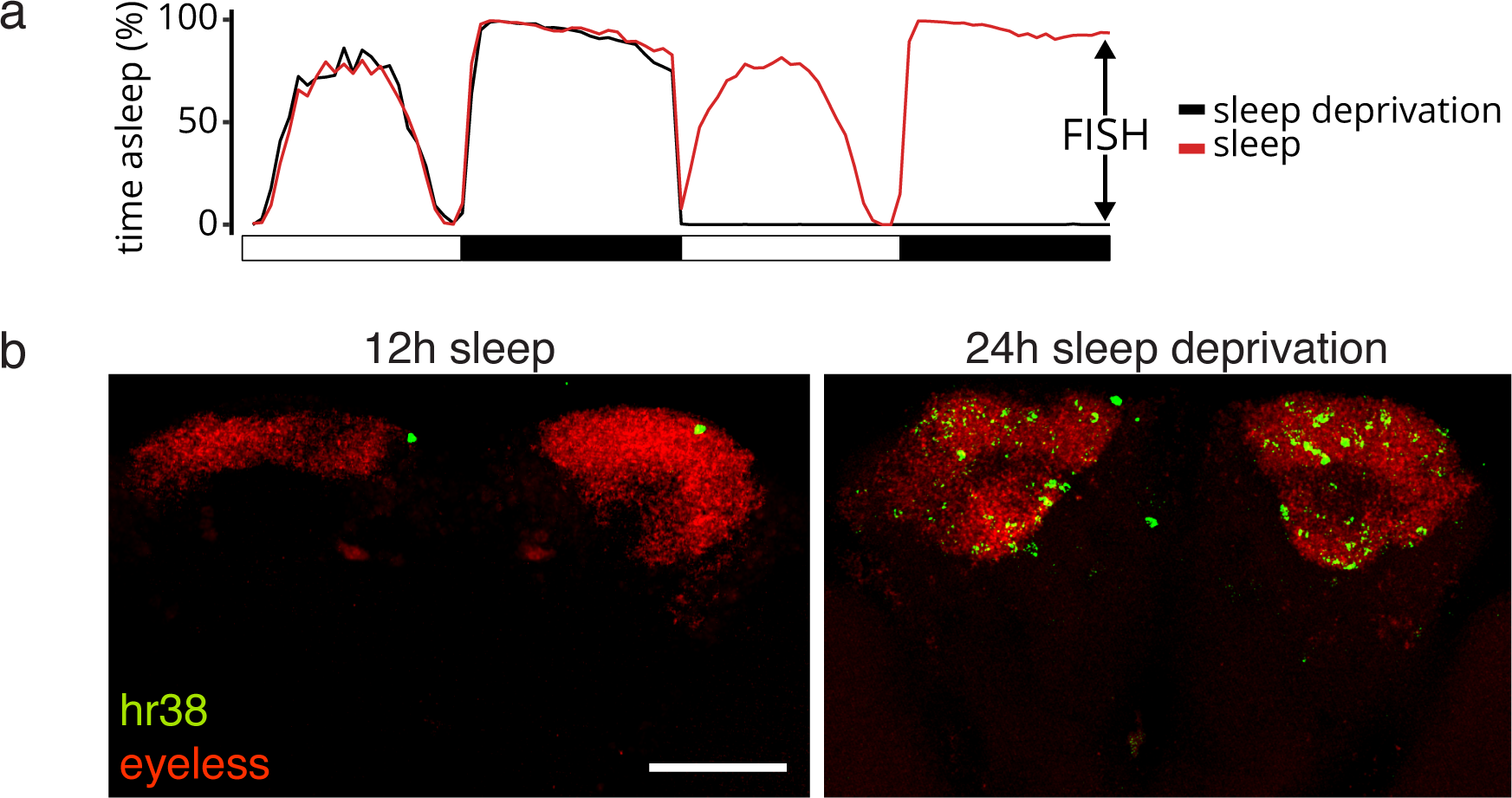
Validation of candidate sleep/wake correlate HR38 in KC. **a.** Sleep and sleep deprivation behaviour traces of flies used for candidate gene validation (sleep: n=256, SD: n=232). Arrows indicate time of sampe collection. **b.** Representative images of validating candidate gene HR38 with KC marker gene eyeless by fluorescent in situ hybridisation. Scale bar, 50 µm.

**Extended Data Figure 9.**
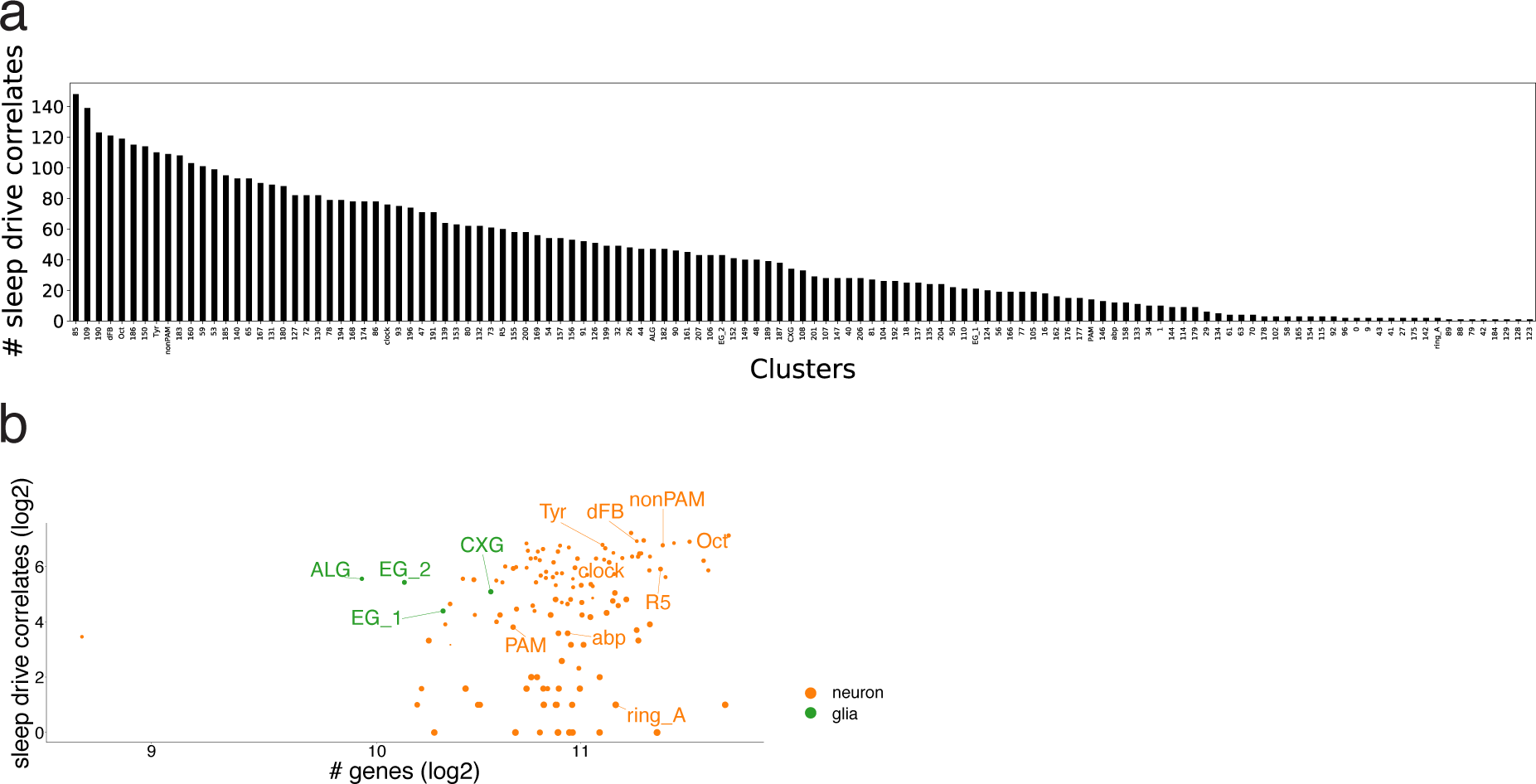
Sleep drive correlates across all clusters. **a.** Number of sleep drive correlates across all clusters. **b.** Correlation between number of genes and number of sleep drive correlates for neuronal and glial cell types.

**Extended Data Figure 10.**
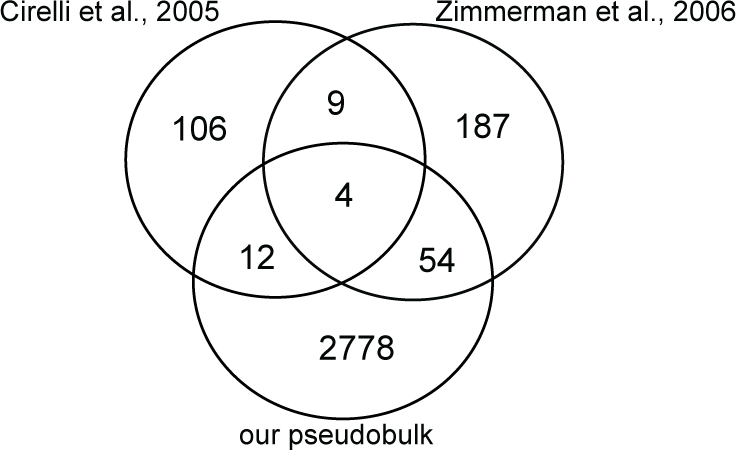
Bulk transcriptomics is insufficient to capture molecular sleep/wake correlates. Comparing sleep/wakefulness correlates across all cells combined found in this study with two previous bulk microarray studies (Cirelli et al. 2005, Zimmerman et al. 2006), that similarly profiled differences between sleep and wakefulness conditions, shows that the overlap of significant genes is minimal between all three studies.

**Supplementary Table 1.** Overview of used fly lines by run, condition and ZT time.

| Run | DGRP_Line | ZT_Time | Treatment | Condition | Age (days) | number brains |
| --- | --- | --- | --- | --- | --- | --- |
| run 1 | 441 | 20 | Sleep | S ZT 12-20 | 6_9 | 10 |
| run 1 | 379 | 20 | Wakefulness | SD ZT12-20 | 6_7 | 10 |
| run 1 | 88 | 8 | Wakefulness | W ZT0-8 | 6_9 | 10 |
| run 1 | 313 | 8 | Wakefulness | SD ZT0-8 | 6_9 | 6 |
| run 1 | 359 | 8 | Wakefulness | SD ZT0-8 | 6_7 | 4 |
| run 2 | 303 | 20 | Sleep | S ZT 12-20 | 7 | 9 |
| run 2 | 313 | 20 | Sleep | S ZT 12-20 | 6 | 1 |
| run 2 | 88 | 20 | Wakefulness | SD ZT12-20 | 7 | 8 |
| run 2 | 908 | 20 | Wakefulness | SD ZT12-20 | 7 | 2 |
| run 2 | 441 | 2 | Wakefulness | SD ZT12-2 | 7 | 7 |
| run 2 | 892 | 2 | Wakefulness | SD ZT12-2 | 7 | 3 |
| run 2 | 646 | 2 | Sleep | RS ZT0-2 | 6_7 | 4 |
| run 2 | 379 | 2 | Sleep | RS ZT0-2 | 7 | 5 |
| run 2 | 287 | 2 | Sleep | RS ZT0-2 | 7 | 1 |
| run 3 | 441 | 20 | Sleep | YC-S ZT12-20 | 7 | 7 |
| run 3 | 313 | 20 | Sleep | YC-S ZT12-20 | 7 | 4 |
| run 3 | 379 | 20 | Sleep | S ZT 12-20 | 7 | 6 |
| run 3 | 908 | 20 | Sleep | S ZT 12-20 | 7 | 3 |
| run 3 | 88 | 2 | Sleep | RS ZT0-2 | 7 | 2 |
| run 3 | 303 | 2 | Sleep | RS ZT0-2 | 7 | 6 |
| run 3 | 892 | 2 | Sleep | RS ZT0-2 | 6_7 | 2 |
| run 3 | 646 | 2 | Wakefulness | SD ZT12-2 | 6_7 | 4 |
| run 3 | 287 | 2 | Wakefulness | SD ZT12-2 | 7 | 6 |
| run 4 | 441 | 2 | Sleep | RS ZT0-2 | 7 | 6 |
| run 4 | 287 | 2 | Sleep | RS ZT0-2 | 7 | 4 |
| run 4 | 379 | 14 | Sleep | S ZT12-14 | 6_7 | 13 |
| run 4 | 646 | 14 | Sleep | S ZT12-14 | 6 | 2 |
| run 4 | 908 | 14 | Wakefulness | SD ZT12-14 | 6_7 | 6 |
| run 4 | 88 | 14 | Wakefulness | SD ZT12-14 | 6_7 | 4 |
| run 5 | 88 | 20 | Sleep | S ZT 12-20 | 7 | 7 |
| run 5 | 287 | 20 | Sleep | S ZT 12-20 | 7 | 7 |
| run 5 | 379 | 20 | Sleep | S ZT 12-20 | 7 | 7 |
| run 5 | 303 | 20 | Wakefulness | SD ZT12-20 | 6_7 | 7 |
| run 5 | 441 | 20 | Wakefulness | SD ZT12-20 | 6_7 | 7 |
| run 5 | 908 | 20 | Wakefulness | SD ZT12-20 | 7 | 7 |
| run 6 | 287 | 8 | Wakefulness | W ZT0-8 | 6_7 | 7 |
| run 6 | 441 | 8 | Wakefulness | W ZT0-8 | 5_7 | 7 |
| run 6 | 303 | 8 | Wakefulness | SD ZT12-8 | 6_7 | 7 |
| run 6 | 379 | 8 | Wakefulness | SD ZT12-8 | 7 | 7 |
| run 6 | 646 | 8 | Sleep | GBX-S ZT12-8 | 4_6 | 9 |
| run 6 | 908 | 8 | Sleep | GBX-S ZT12-8 | 5_7 | 7 |
| run 7 | 303 | 8 | Wakefulness | W ZT0-8 | 6_7 | 6 |
| run 7 | 379 | 8 | Wakefulness | W ZT0-8 | 7 | 7 |
| run 7 | 646 | 8 | Wakefulness | SD ZT12-8 | 4_7 | 7 |
| run 7 | 908 | 8 | Wakefulness | SD ZT12-8 | 7_8 | 7 |
| run 7 | 287 | 8 | Sleep | GBX-S ZT12-8 | 7 | 7 |
| run 7 | 441 | 8 | Sleep | GBX-S ZT12-8 | 6_8 | 7 |

**Supplementary Table 2.** Cell type annotations.

| annotation | resolution | cluster |
| --- | --- | --- |
| CXG | 2 | 76 |
| ALG | 8 | 19 |
| ALG_0 | 8 | 133 |
| PG | 4 | 81 |
| EG_1 | 4 | 28 |
| EG_2 | 8 | 64 |
| EG_0 | 2 | 18 |
| ab | 2 | 4 |
| abp | 2 | 19 |
| y | 3 | 0 |
| nonPAM | 8 | 136 |
| PAM | 8 | 37 |
| clock | 8 | 84 |
| Tyr | 3 | 105 |
| Oct | 3 | 103 |
| 5-HT | 8 | 151 |
| ring_2 | 8 | 95 |
| ring_1 | 8 | 100 |
| adPN | 3 | 31 |
| PB | 2 | 43 |
| T4/T5 | 2 | 38 |
| Tm1/TmY8 | 3 | 101 |
| Mi1 | 3 | 91 |
| T1 | 8 | 135 |
| ring_A | part of ring_2 (except for R5) |  |
| ring_B | top part of ring_2 and ring_1 |  |
| ring_C | R5, part of ring_2 |  |
| dFB | 8 | 198 |

**Supplementary Table 3.**
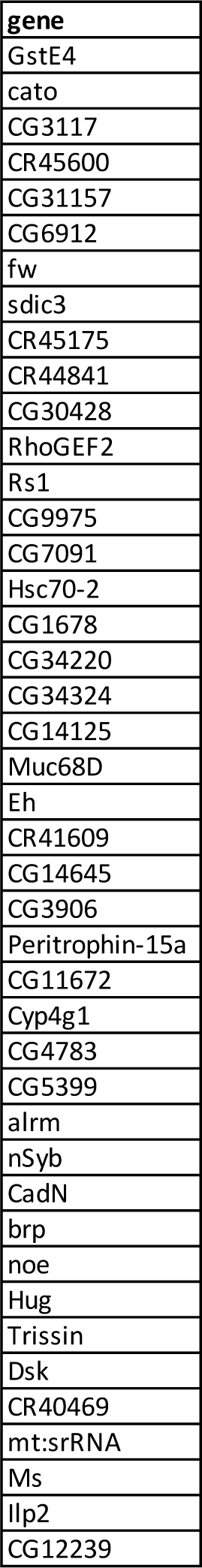
Batch effect genes between runs or genotypes.

**Supplementary Table 4.** Statistics by run.

| Sample | Reads | Sequencing Saturation (%) | Cells pre-filter | Median Genes | Median UMIs | Singlets (demuxlet) | Mitochondrial % Threshold |
| --- | --- | --- | --- | --- | --- | --- | --- |
| DGRP_Mix_Sleep_Wakefulness_Run1 | 517,406,996 | 60.1 | 22358 | 1604 | 4334 | 18856 | 20 |
| DGRP_Mix_Sleep_Wakefulness_Run2 | 1,324,890,675 | 59.3 | 36984 | 1829 | 6333 | 22225 | 30 |
| DGRP_Mix_Sleep_Wakefulness_Run3 | 1,422,773,423 | 59.9 | 31872 | 1930 | 6902 | 22769 | 20 |
| DGRP_Mix_Sleep_Wakefulness_Run4 | 1,180,889,558 | 59.3 | 34043 | 1776 | 5920 | 16594 | 30 |
| DGRP_Mix_Sleep_Wakefulness_Run5 | 304,007,531 | 56.1 | 15130 | 1581 | 4655 | 11810 | 25 |
| DGRP_Mix_Sleep_Wakefulness_Run6 | 342,832,941 | 57.2 | 13410 | 1534 | 4334 | 13543 | 25 |
| DGRP_Mix_Sleep_Wakefulness_Run7 | 211,110,425 | 59.4 | 10241 | 1393 | 3661 | 9383 | 25 |

